# Longitudinal analysis of retinal cell state transitions in *RB1*-deficient retinal organoids reveals the nascent cone precursors are the earliest cell-origin of human retinoblastoma

**DOI:** 10.1101/2025.03.13.642537

**Authors:** Ke Ye, Yuan Wang, Ping Xu, Bingbing Xie, Shijing Wu, Wenxin Zhang, Guanjie Gao, Dandan Zheng, Xiaojing Song, Suai Zhang, Fuying Guo, Yongping Li, Yizhi Liu, Jie Wang, Ruifang Sui, Xiufeng Zhong

## Abstract

All cancers develop from malignant transformation of normal cells, while it is challenging to identify these “cells-of-origin” since it is impossible to witness the dynamic changes in human cancers. Retinoblastoma (Rb), an intraocular malignant cancer, is a typical model for study the molecular and cellular mechanisms of cancers. Although the maturing cone precursors (CPs) are proposed as the cellular origin of human Rb, it is unknow whether other retinal cells are also sensitive to *RB1* inactivation. Here we developed *RB1*-deficient human retinal organoids (ROs) models from *RB1*^−/−^ or *RB1*^+/−^ human induced pluripotent stem cells (hiPSCs). *RB1*^−/−^ hiPSCs developed into Rb tumors which recapitulated the features of human native Rb and had the ability to form consecutive orthotopic xenografts in mice. Importantly, *RB1* loss induced the overproliferation of ATOH7^+^ neurogenic retinal progenitor cells (nRPCs), which disrupted the retinal development by generating proliferative, nascent retinal cells including retinal ganglion cells and CPs. scRNA-seq analysis verified the ATOH7^+^/RXRγ^+^ nascent CPs survived and finally drove Rb development. In contrast, monoallelic *RB1* inactivation with low pRB expression did not induce nascent CPs proliferation, but only induced nRPCs overproliferation which caused retinocytoma-like phenotype. Finally, a potential therapeutic agent for Rb was identified from multi-omics data. Our findings firstly indicate that nRPCs are the most sensitive cells to *RB1* loss inducing nascent retinal cells abnormal proliferation, and ATOH7^+^ nascent CPs are the earliest cellular origin of human Rb, facilitating drug development for Rb.

## Introduction

Retinoblastoma (Rb) is a pediatric eye cancer that originates in the developing retina and is usually diagnosed before the age of five^1^. Most Rb is caused by biallelic inactivation of the *RB1* gene, the first identified tumor suppressor gene^2^. It encodes pRB, which controls the G1/S transition of the cell cycle by modulating the activity of E2F transcription factors^3^. Therefore, pRB plays an important role in tumorigenesis by regulating cell-cycle progression and cell differentiation^4,5^. Because pRB is expressed in cell-type specific and stage-dependent manner during retinogenesis^6^, the complex expression pattern makes the understanding of pRB biology in human retina is still incomplete, including when or how tumorigenesis begins.

Cellular origin of Rb is a most challenging and significant question for long term. Previous studies with human Rb samples suggest that Rb may arise from retinal progenitor cells (RPCs), photoreceptor cells (PRCs) and interneurons since Rb display high heterogeneity at the molecular and cellular level^7–11^. However, because most samples are collected in the late stages of tumor development where cell types likely cannot represent the original cells, which are normal cells acquiring caner-initiating mutations leading to frank tumor growth, it is difficult to identify when or how tumorigenesis begins. Studies with human fetal retina or explants by shRNA mediated *RB1* depletion propose that ARR3^+^ matured cone precursors (CPs) are the potential cell-of-origin of human Rb^9,10^. However, it is still unknow whether other retinal cells are also sensitive to *RB1* inactivation to drive tumor development. In addition, mouse models with *RB1* loss does not recapitulate the features of human Rb due to species variation^8,12,13^. There is a need to further clarify these fundamental questions by using a novel, accessible and reproducible human retinogenesis model: retinal organoids (ROs)^14^.

ROs are induced from pluripotent stem cells and are capable of recapitulating the entire process of retinogenesis *in vitro*^15^. During ROs differentiation, all seven major retinal cell subtypes including retinal ganglion cells (RGCs), cones, horizontal cells (HCs), amacrine cells (ACs), muller glia cells (MGs), bipolar cells (BCs) and rods develop from RPCs in a time-dependent manner^15,16^. Several studies have attempted to model human Rb using RO models, but not all studies succeed. Zheng et al. reported that *RB1*-deleted ROs did not induce Rb formation *in vitro* and *in vivo*, but only caused the increasing apoptosis^17^. Norrie et al. developed Rb orthotopic xenografts by injecting *RB1* mutated ROs-derived cells at day 45, and proposed Rb tumors should be progenitor signature cells^18^. Jin’s group observed Rb formation in *RB1*-null ROs at day 120 from which orthotopic xenografts in mice acquired, and suggest matured CPs are cellular origin of Rb^19^. Similarly, Rozanska et al. also observed a similar cell-of-origin to Jin’s research^20^. However, a recent study raises a possibility that the early CPs exhibited a potential effect in Rb initiation^21^. Therefore, a detailed analysis is necessary to discriminate the specific state and explore how these cells contribute to Rb development. In this study, we established pRB-negative and pRB-decreased RO models from *RB1*^−/−^ or *RB1*^+/−^ hiPSCs and deciphered malignant transformation of normal retinal cells during the whole process of retinogenesis. Only ROs with complete loss of *RB1* developed Rb tumors mimicking the features of human native Rb with ability to form consecutive orthotopic xenografts in mice. ATOH7^+^ nRPCs are the most sensitive cells to *RB1* loss inducing nascent retinal cells abnormal proliferation, and ATOH7^+^/RXRγ+ nascent CPs are the earliest cellular origin of human Rb. In contrast, the monoallelic *RB1* inactivation with low pRB expression did not induce nascent CPs abnormal proliferation, but induced nRPCs overproliferation leading to a retinocytoma-like lesion due to the enrichment of non-proliferative CPs. Multi-omics analysis of *RB1*^−/−^ ROs identified a potential therapeutic agent for Rb. Our findings provide new insights into cellular and molecular mechanisms of human Rb and facilitate therapeutic development.

## Results

### 1. *RB1* deletion induces tumor formation of Rb in ROs

Two clones of *RB1*^−/−^ hiPSC lines (C1 and C5) were generated using CRISPR/Cas9 technology (Fig. S1A, B). *RB1*^−/−^ hiPSC lines exhibited downregulation of pRB protein level without affecting the typical features of hiPSCs (Fig. S1C-H). We did not identify differences in both the cell cycle and the potential to form teratomas between WT and *RB1*^−/−^ hiPSCs (Fig. S2A-C). To generate *RB1*-inactivated hiPSCs under different genetic backgrounds, another KO strategy was performed to generated *RB1*-inactivated hiPSCs from the another hiPSC line (Fig. S3A). One *RB1* biallelic-deleted hiPSC line (*RB1*^Puro/Puro^) and one *RB1* heterozygous-deleted hiPSC line (*RB1*^W/Puro^) were obtained (Fig. S3B-F).

RO differentiation was performed with our optimized protocols (Fig. 1A)^15,22^. No significant disorders were observed from embryoid body to early *RB1*^−/−^ ROs (Fig. S4A). pRB^+^ cells were only detected in the WT ROs (Fig. S4B). After 70 days of differentiation, *RB1*^−/−^ ROs became larger in size than WT ROs (Fig. 1B, C), and the ratio of Ki67^+^ and EdU^+^ cells increased (Fig. 1D, E, S4C, D). Rb markers, including SYK and p16^INK4a^ were only detected in *RB1*^−/−^ ROs on day 90 (Fig. 1F, G, S4E). Transcriptomic analysis revealed the upregulation of cell cycle-(*MKI67*, *CCNE2* etc.) and cancer-related genes (*SYK*, *MCM2* etc.) in *RB1*^−/−^ ROs (Fig. 1H), and GSEA confirmed the enrichment of Rb-related genes (Fig. S4G). Morphologically, *RB1*^−/−^ ROs lost laminated structure, comprised homogenous tumor-like cells with a larger nuclear size and mitotic figures (Fig. 1I, J, S4F). IF analysis showed that the majority of cell population in *RB1*^−/−^ ROs expressed CPs marker RXRγ, which is similar to clinical Rb samples (Fig. 1K, L). The same phenomenon was identified in other *RB1* homozygous mutated cell lines (Fig. S4H, I and S5). Taken together, these results demonstrate that *RB1* loss can robustly induce tumorigenesis of cone-rich Rb in RO model.

**Fig. 1.**
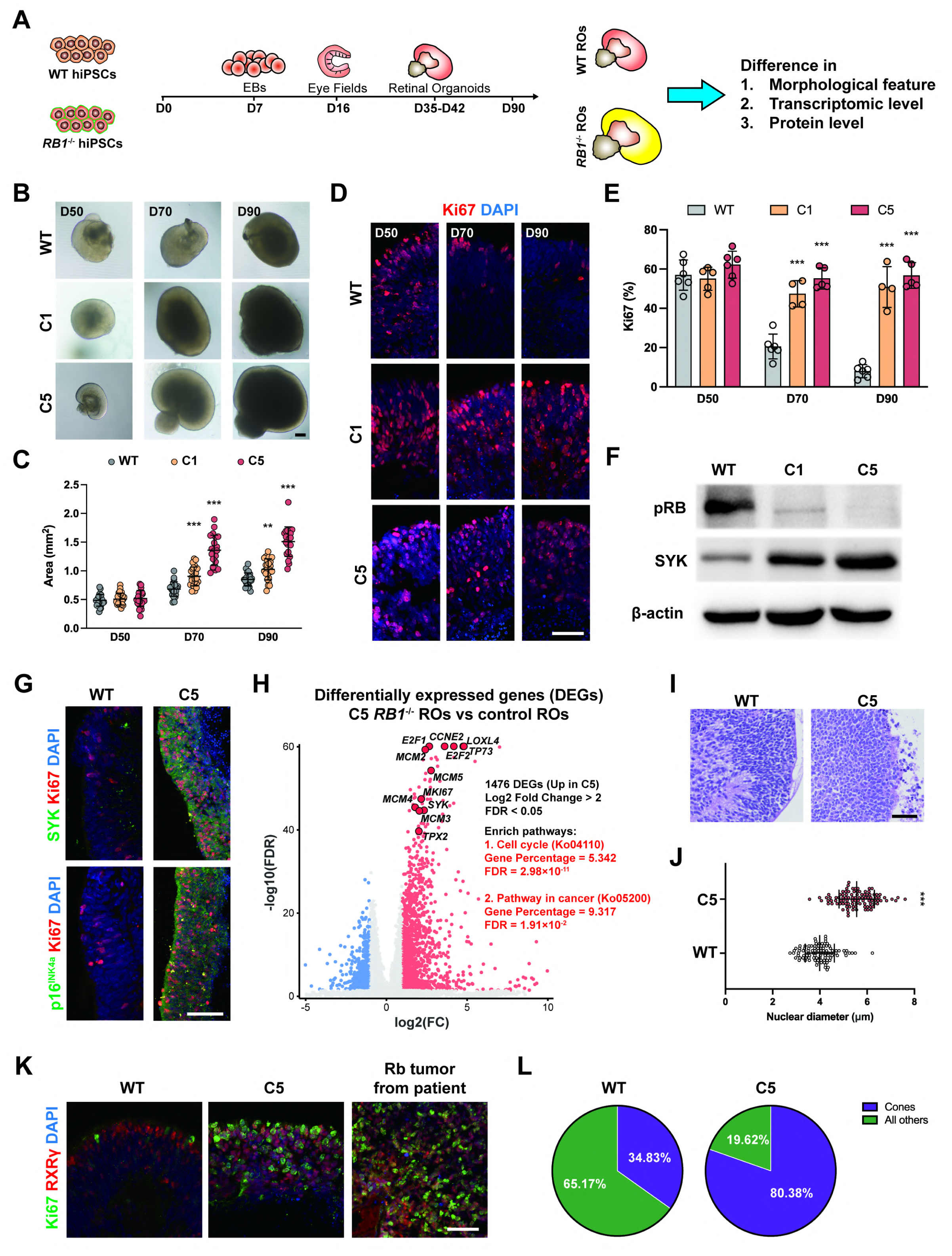
Tumorigenesis of ROs with *RB1* loss *in vitro*. (A) Target strategy of analysis of *RB1*^−/−^ ROs. (B) Representative microscopic images of WT and *RB1*^−/−^ ROs from day 50 to day 90. (C) Quantification of size of ROs in different culture phase. Data represent mean ± SD. *** P < 0.001 vs. WT, n ≥ 20. (D) Representative immunostaining for Ki67 in WT and *RB1*^−/−^ ROs from day 50 to 90. (E) Quantification of Ki67^+^ cells from day 50 to 90. Data represent mean ± SD. *** P < 0.001 vs. WT, n = 4-6. (F) Western blot analysis of pRB and SYK protein level in WT and *RB1*^−/−^ ROs at day 90. (G) Representative immunostaining for SYK, p16^INK4a^ and Ki67 in WT and *RB1*^−/−^ ROs at day 90. (H) Volcano plot visualizations of DEGs and related enrich pathways from bulk RNA-seq data obtained between WT and C5 *RB1*^−/−^ ROs. (I) Representative H&E staining of WT and C5 *RB1*^−/−^ ROs at day 80. (J) Quantification of nuclear size of WT and C5 *RB1*^−/−^ ROs. Data represent mean ± SD. *** P < 0.001 vs. WT, n ≥ 20. (K) Representative immunostaining for Ki67 and RXRγ in WT, C5 *RB1*^−/−^ ROs and patient tumor sample. (L) Quantification of RXRγ^+^ cells in WT and C5 *RB1*^−/−^ ROs. Scale bars = 200 μm (B) and 50 μm (D, G, I, K).

### 2. Formation of consecutive Rb orthotopic xenografts from *RB1*^−/−^ ROs in immunodeficient mice

Since xenograft tumor formation are the golden standard to confirm malignant feature of cells, we investigated whether retinal tumor-like cells in *RB1*^−/−^ ROs recapitulated the formation and characteristics of human Rb in mice eyes^23^. The tumor-like cells from D70 *RB1*^−/−^ ROs were engrafted into the intravitreal or subretinal spaces of NOD/SCID mice to generate primary xenograft tumors (Fig. 2A). 10 weeks after injection (WAI), SD-OCT identified highly reflective tumor-like masses in some mice (Fig. 2B). 13 WAI, anterior chambers of mice eyes were full of whitish tumor cells (Fig. 2C). After enucleation, tumor cells were identified in 7 out of 10 mice in *RB1*^−/−^ group (Fig. 2D, table S1). Hematoxylin and eosin (H&E) staining showed the tumor cells formed numerous F-W rosettes with mitosis figures, which were similar to well-differentiated human Rb (Fig. 2E, S6A). Transmission electron microscopy (TEM) analysis confirmed that the tumor cells exhibited Flexner-Wintersteiner rosettes, high nucleocytoplasmic ratio and abundant mitochondria (Fig. S6B). In contrast, we did not observe any tumors in WT group (Fig. S6C-E).

**Fig. 2.**
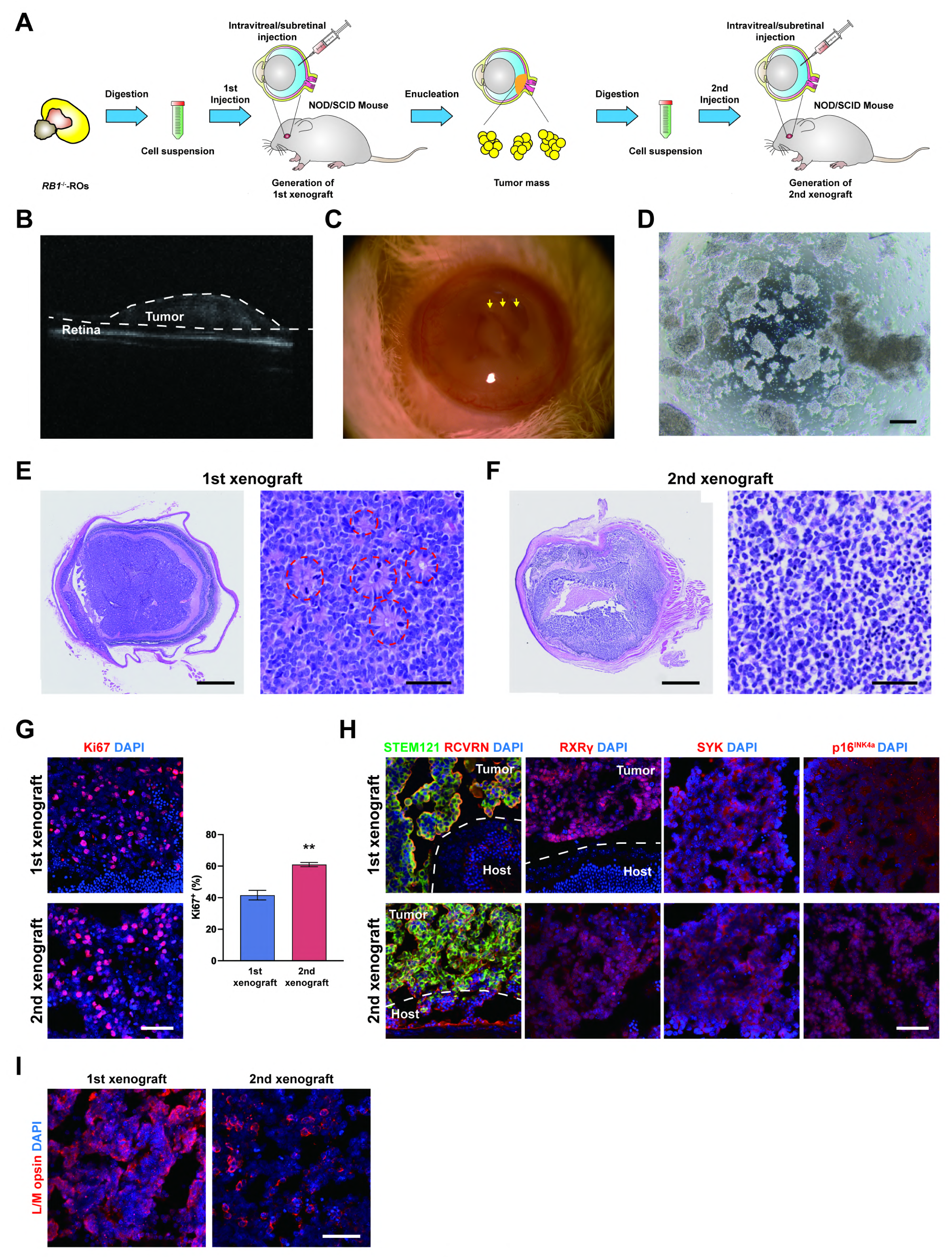
Xenograft of ROs-derived cells into mice *in vivo*. (A) Schematic outline of the generation of 1st and 2nd xenograft. (B) Representative optical coherence tomography (OCT) images of engrafted eyes at 10 WAI from *RB1*^−/−^ ROs. An obvious highly reflective tumor was observed. (C) Representative slit-lamp images of engrafted eyes at 13 WAI from *RB1*^−/−^ ROs. Whitish tumor cells were observed in anterior chambers. (D) Representative microscopic image of tumor mass derived from 1st xenograft. (E) Representative H&E staining images of 1st xenograft from *RB1*^−/−^ ROs. The dash lines indicate the rosette structure. (F) Representative H&E staining images of 2nd xenograft with injection of 1st tumor cells. No rosettes were observed. (G) Representative immunostaining for Ki67 and quantification of Ki67^+^ cells in 1st and 2nd xenograft. ** P < 0.01 vs. 1st xenograft, n = 3. (H) Representative immunostaining for human specific marker (STEM121), cone markers (RCVRN, RXRγ) and Rb markers (SYK, p16^INK4a^) in 1st and 2nd tumor. (I) Representative immunostaining for L/M opsin in 1st and 2nd tumor. Scale bars = 200 μm (D), 500 (left), 50 (right) μm (E, F) and 50 μm (G, H, I).

To further explore the tumorigenicity, cells isolated from primary xenograft were injected into 5 new mice. All mice developed eye tumors 13 WAI (Table S2). Surprisingly, the 2nd xenograft tumors did not have visible rosette structures, but presented necrosis region (Fig. 2F), suggesting tumor growth quite fast. IF analysis confirmed the higher ratio of Ki67^+^ cells in the 2nd xenograft (Fig. 2G). These features were more similar to poorly-differentiated human Rb or xenograft tumors from Rb cell lines (Fig. S6A, F). Both of the 1st and 2nd xenograft tumors expressed cone markers and Rb markers (Fig. 2H), but did not express markers specific for other retinal cell types (Fig. S6G). L/M opsin highly expressed in 1st xenograft but low in the 2nd xenograft, consistent with their morphologic features, high and low differentiated Rb tumors respectively (Fig. 2I). These results demonstrate that retinal tumor-like cells from *RB1*^−/−^ ROs have ability to develop into the typical phenotype of Rb tumors with strong expansion capacity.

### 3. *RB1* loss induced overproliferation of nRPCs generating proliferative nascent retinal cells in *RB1*^−/−^ ROs

Here we explore how *RB1* deletion affects the retinal development in ROs. During early retinogenesis, RPCs gradually develop into ATOH7^+^ nRPCs, which generate CPs and RGCs (Fig. 3A)^24^. In WT ROs, ATOH7^+^ cells were mostly located in neuroblast layer (NBL) and ganglion cell layer (GCL) (Fig. 3B). In *RB1*^−/−^ ROs, the proliferative cells among ATOH7^+^ cells was significantly higher than that in WT ROs by counting Ki67^+^ and EdU^+^ cells with disorder distribution across all layers (Fig. 3C-E, S7A-D). It induced more ATOH7^+^ cells in *RB1*^−/−^ ROs from day 35 (Fig. S7E). In nRPCs-derived offsprings, a higher portion of ATOH7^+^ cells co-expressed with RXRγ or HuD in *RB1*^−/−^ ROs (Fig. 3F), implying that more CPs and RGCs were generated. Notably, the postmitotic CPs and RGCs did not express Ki67 in WT ROs, but did in *RB1*^−/−^ ROs (Fig. 3G, H). RNA-seq and IF analysis revealed that the expression of *RB1*-targetted genes were significantly increased in *RB1*^−/−^ ROs from day 50 (Fig. 3I, S7F), which revealed that pRB loss induced the cell cycle regulation in early retinogenesis.

**Fig. 3.**
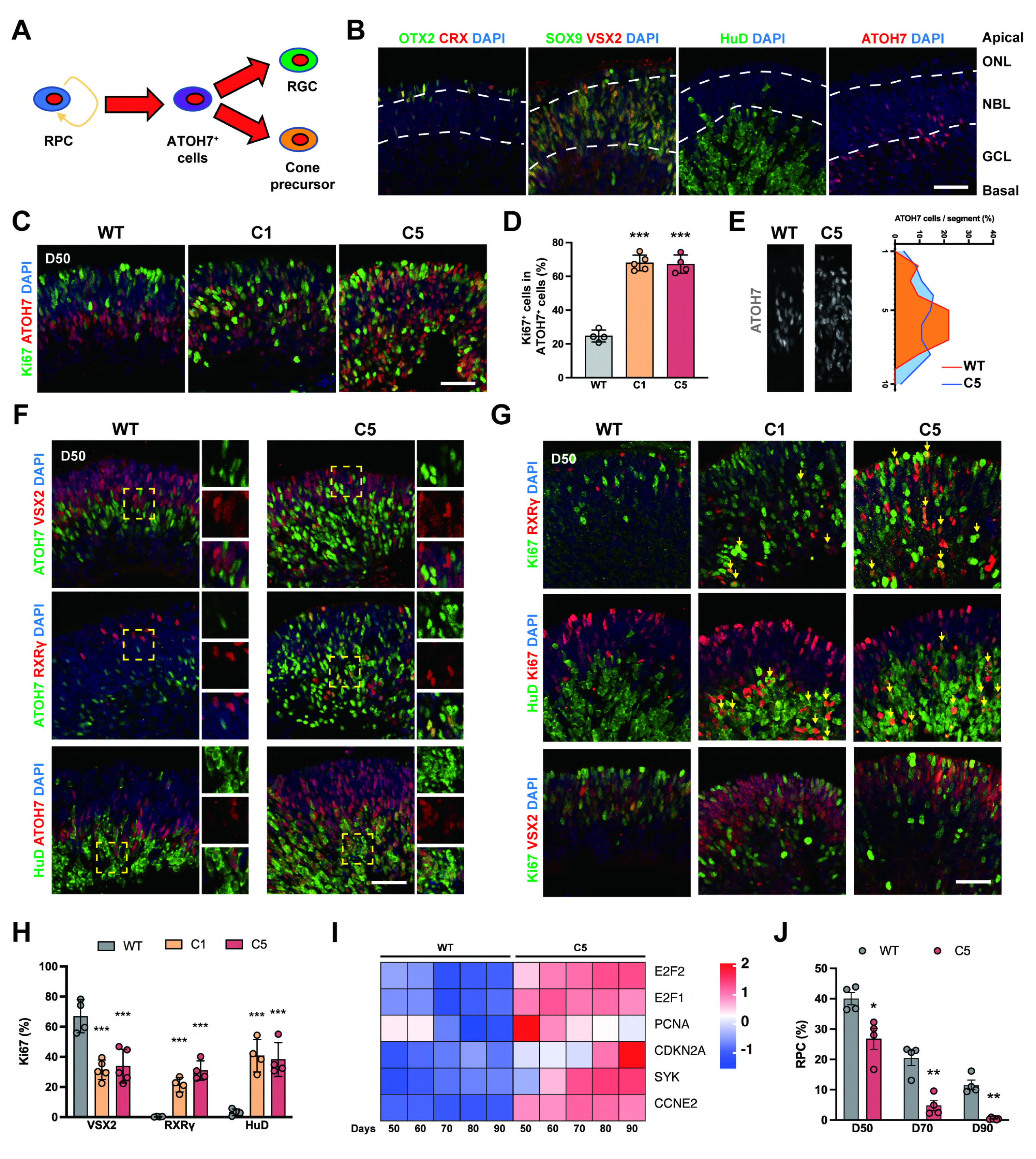
Developmental defect of late retinogenesis due to pRB expression dependent nRPCs proliferation. (A) Summary of the differentiation of early-born retinal cells from ATOH7^+^ cells. (B) Representative immunostaining for early PRC markers (OTX2, CRX), RPC markers (VSX2, SOX9), RGC marker (HuD) and ATOH7 in WT ROs showing the localization of each type of cells. ONL: outer nuclear layer; NBL: neuroblastic cell layer; GCL: ganglion cell layer. (C) Representative immunostaining for Ki67 and ATOH7 in WT and *RB1*^−/−^ ROs at day 50. (D) Quantification of Ki67^+^ cells in ATOH7^+^ cells in WT and *RB1*^−/−^ ROs at day 50. Data represent mean ± SD. *** P < 0.001 vs. WT, n = 4-5. (E) Representative immunostaining for ATOH7 and relative distribution of ATOH7^+^ cells in WT and *RB1*^−/−^ ROs at day 50. (F) Representative immunostaining for VSX2, RXRγ, HuD and ATOH7 in WT and *RB1*^−/−^ ROs at day 50. (G) Representative immunostaining for VSX2^+^, RXRγ^+^, HuD^+^ and Ki67 in WT and *RB1*^−/−^ ROs at day 50. Arrows indicate the Ki67^+^ cells in different cell types. (H) Quantification of Ki67^+^ cells in RXRγ^+^, HuD^+^ and VSX2^+^ cells in WT and *RB1*^−/−^ ROs at day 50. Data represent mean ± SD. *** P < 0.001 vs. WT, n = 4-5. (I) Heatmap of differential expression of Rb marker genes in WT and *RB1*^−/−^ ROs from day 50 to 90. (J) Cell ratio of RPC from day 50 to day 90. Data represent mean ± SD. * P < 0.05, ** P < 0.01 vs. WT. Scale bars = 50 μm (B, C, F, G).

In contrast, the percentage of Ki67^+^/VSX2^+^ RPCs was rapidly decreased at day 50 (Fig. 3G, H). EdU-labeling confirmed this observation (Fig. S7G, H). This led to a rapid decrease in the progenitor pool in *RB1*^−/−^ ROs in terms of cell ratio and mRNA level (Fig. 3J, S7I, J). These results suggested that *RB1* loss not only induced the proliferation of nRPC-generating nascent CPs and RGCs, but also potentially caused RPC differentiation defect.

### 4. Longitudinal analysis showed that pRB loss induced CPs tumor-like clonal growth at the expense of specification of late born retinal cells

During maturation of *RB1*^−/−^ ROs, the expression of retinal cell-specific genes other than cones were downregulated, including RGC-specific genes (Fig. 4A). We found that *RB1* loss also indued cell apoptosis from day 70 by analysis of cleaved caspase-3 (CC3)^+^ cells (Fig. S8A, B). pRB-negative RGCs were more susceptible to apoptosis than CPs (Fig. S8C, D). IF analysis for multiple retinal markers at day 90 also confirmed that the pRB-negative ROs lost the ability to develop into other retinal cell types except cones presenting tumor-like clonal growth (Fig. 4B, S9A, B).

**Fig. 4.**
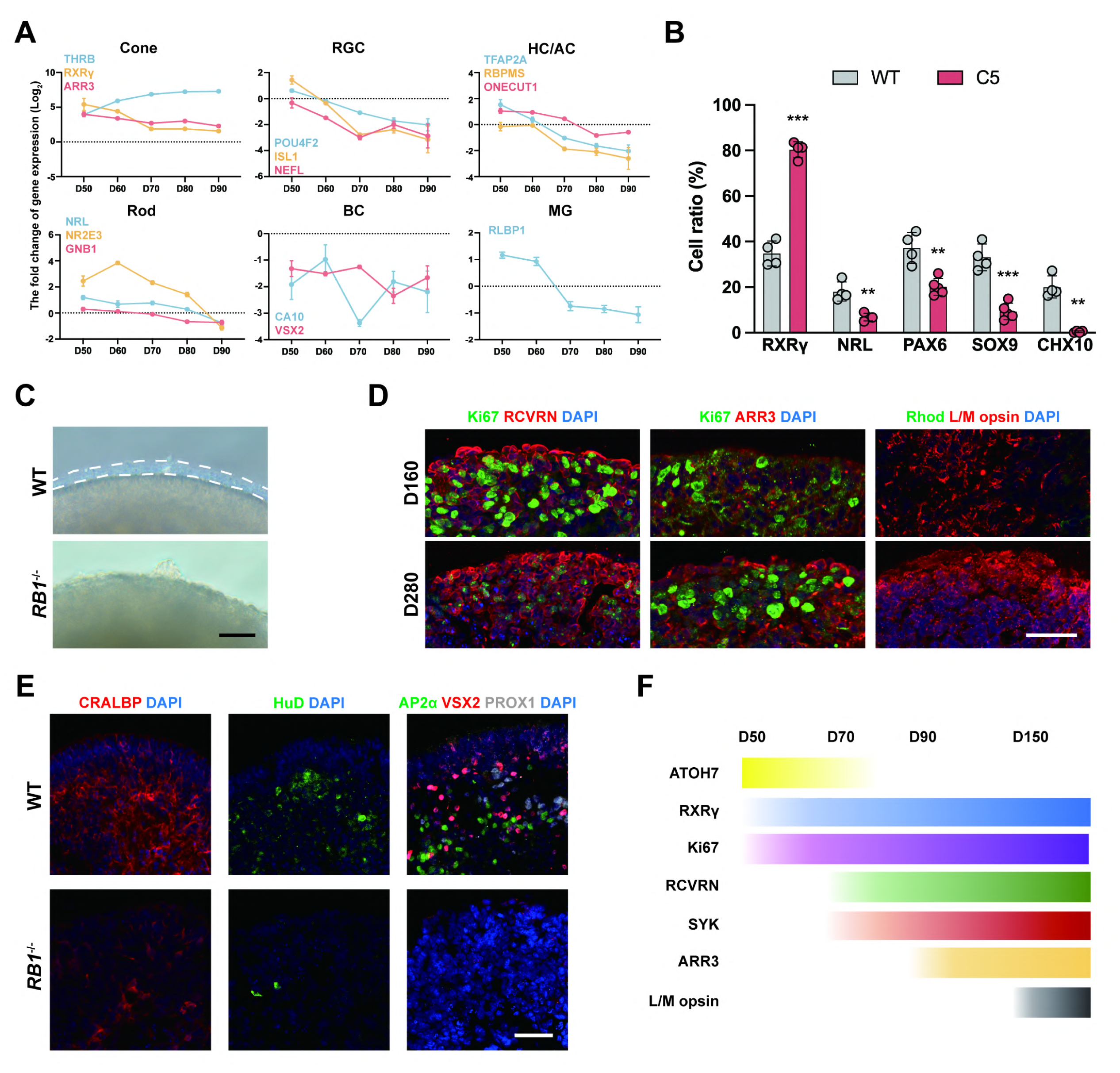
Longitudinal analysis of pRB-negative Ros. (A) The fold change of expression of cell type specific marker genes in WT and *RB1*^−/−^ ROs from day 50 to 90, n = 3. (B) Quantification of the ratio of RXRγ^+^, NRL^+^, SOX9^+^, PAX6^+^ and VSX2^+^ cells in WT and *RB1*^−/−^ ROs at day 90. Data represent mean ± SD. ** P < 0.01, *** P < 0.001 vs. WT, n = 4-5. (C) Representative microscopic images of WT and *RB1*^−/−^ ROs at day 160. (D) Representative immunostaining for RCVRN, ARR3, Ki67, L/M opsin and Rhodopsin in *RB1*^−/−^ ROs at day 160 and 280. (E) Representative immunostaining for CRALBP^+^ (MGs), HuD^+^ (RGCs), AP2α^+^ (HCs), PROX1^+^ (ACs), and VSX2^+^ (BCs) in WT and RB1-/- ROs at day 280. (F) Summary of temporal expression of nRPC, cone and Rb markers during tumor development. Scale bars = 50 μm (D, E) and 200 μm (C).

To observe the tumor progression for longer time, *RB1*^−/−^ ROs were cultured up to day 280. *RB1*^−/−^ ROs displayed a disorganized morphology without outer segments formation (Fig. 4C). In the WT ROs, RCVRN^+^ photoreceptor precursors grew into mature photoreceptors that expressed L/M opsin and rhodopsin (Fig. S9C)^15,25,26^. Interestingly, *RB1*-deleted ROs also expressed the L/M opsin in the same phase with SYK and Ki67 expression (Fig. 4D, S9D). However, rhodopsin^+^ rods were not detected (Fig. 4D). In addition, the population of other retinal cells were hardly identified in *RB1*^−/−^ ROs (Fig. 4E). This cellular architecture was consistent with that of Rb tumor samples with cone-tumor cell maturation at certain degree. In conclusion, the above results demonstrate that pRB loss induced the proliferation of death resistant CPs, which contributed to Rb tumor progression at the expense of the specification of late born retinal cells (Fig. 4F).

### 5. Single-cell transcriptomic profiling reveals nascent CPs are the earliest source of Rb tumor in *RB1*^−/−^ RO models

scRNA-seq was performed to analyze the cellular composition and developmental trajectory of Rb in *RB1*^−/−^ ROs. Before analysis, the cell cycle genes were removed to prevent bias. According to the expression of marker genes, the cells were classified into specific cell types, including RPCs (*VSX2*, *CCND1*), nascent RGCs (*ATOH7*/*POU4F2*), RGCs (low *ATOH7*/high *POU4F2*), nascent pan PRC precursor (*ATOH7*/*OTX2,* HCs/ACs (*TFAP2A*), rods (*NRL*), and nascent cones (*ATOH7*/*RXRG*) (Fig. 5A, B, E, S10A). The loss of RPC pool in *RB1*^−/−^ ROs was identified by the decreasing cell number of RPCs (Fig. 5B, C). Notably, three additional cell populations were found in *RB1*^−/−^ ROs, including a subtype of proliferative ATOH7^+^ cells. This population also highly expressed cone-determination genes including *Olig2* and *Onecut1*^27^, we defined these cells as proliferating nascent CP (P-CP) (Fig. 5A-C, E, S10B, C). Such population of cells was also identified by IF using triple labeling from day 50 (Fig. 5F, G). Because ATOH7 only expressed in the committed stage of CPs^24,28^, these results suggested that the nascent CPs, transient cells, were sensitive to *RB1* inactivation.

**Fig. 5.**
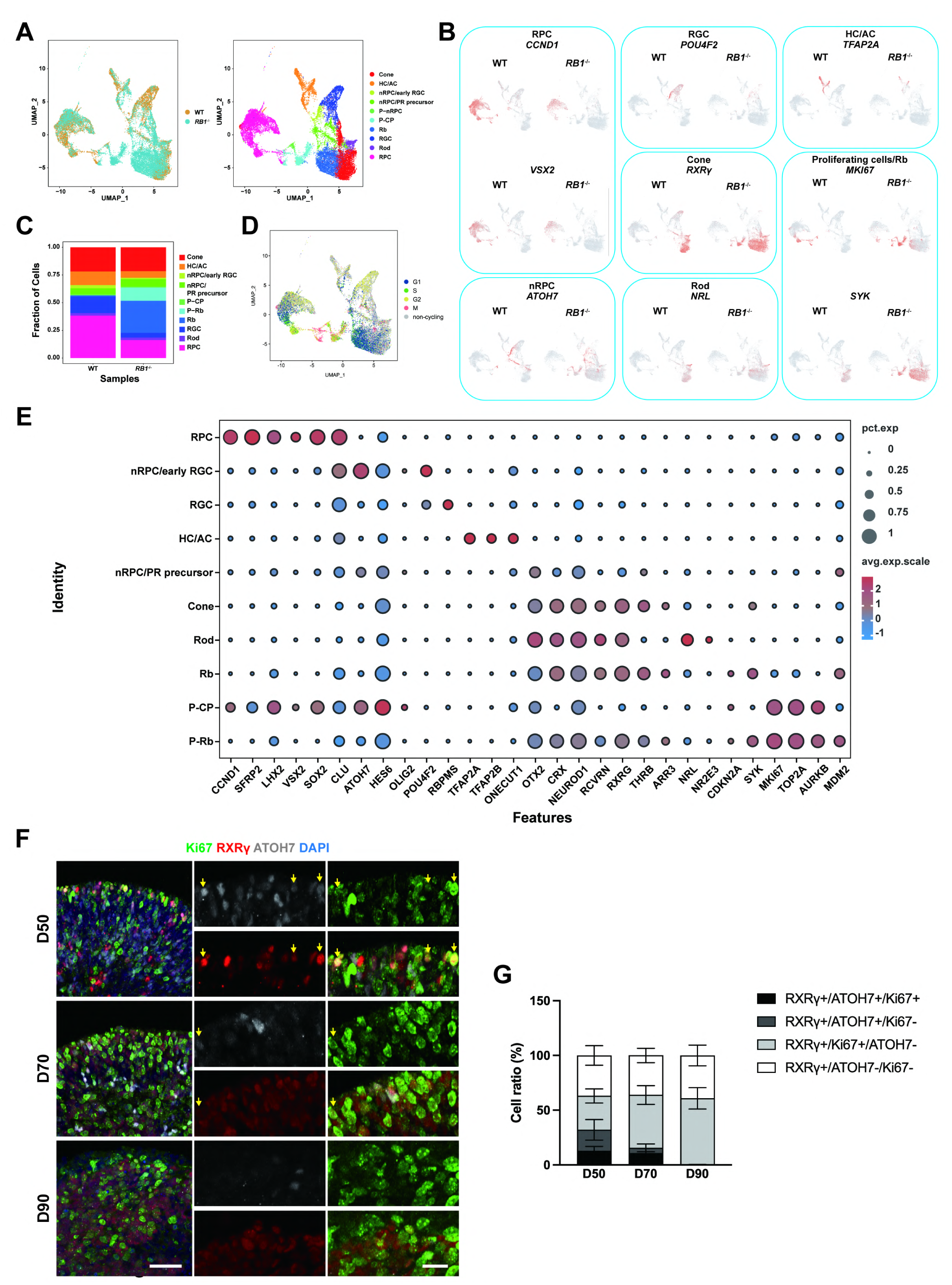
Nascent CPs as a cell source of Rb in scRNA-seq analysis. (A) UMAP plot of unsupervised cell cluster of WT and *RB1*^−/−^ ROs at day 80. Different colors correspond to different cell types. (B) Feature plots of expression of different retinal cell types or Rb cell type-specific markers. (C) Quantification of cell ratio in different cell clusters. (D) Distribution of cells in different cell cycle. (E) Dot plot visualization of expression of different cell type-specific markers. (F) Representative immunostaining for Ki67, RXRγ and ATOH7 in *RB1*^−/−^ ROs from day 50 to day 90. Arrows indicate triple positive cells at different culture day. (G) Quantification of ATOH7^+^ and Ki67^+^ cells in RXRγ^+^ cells in *RB1*^−/−^ ROs from day 50 to day 90. Scale bars = 50 μm (F).

Additionally, one other population highly expressed cone and Rb-related genes (*SYK*, *CDKN2A*, *MDM2*) with lower level of *MKI67*, and was considered as Rb cells (Rb). The proliferative Rb cells (P-Rb) expressed higher levels of *MKI67* and Rb-related genes (*SYK*, *TOP2A*, *MDM2*) (Fig. 5B, C and E, S10B). Cell cycle analysis confirmed that these two additional cell populations re-entered cell cycle (Fig. 5D, S10C). KEGG enrichment analysis of DEGs between Rb-related cell clusters (Rb and p-Rb) and other retinal cells revealed that the former highly expressed cell cycle- and tumor-related genes (Fig. S10D).

Next, we explored how the P-CPs in *RB1*^−/−^ ROs developed into Rb using pseudo-time trajectory analysis by Monocle 2. The data showed that most RPCs presented at the beginning of trajectory and P-CPs mostly present at the end of the trajectory near P-Rb (Fig. S11A, B). These results suggested that these cells may develop directly into proliferative Rb cells. The pseudo-time heatmap showed the progressive dynamic expression of selected Rb genes and cell cycle-related genes (Fig. S11C). Taken together, these data confirm that malignant tumor cells arise from the committed or nascent CPs.

### 6. Low pRB expression did not induce abnormal proliferation of nascent CPs, but nRPCs causing retinocytoma-like phenotype in *RB1*^+/−^ ROs

Two heterozygous mutated hiPSC lines were used to confirm the necessity of biallelic loss of *RB1* in Rb initiation. One hiPSC line (*RB1*^W/Puro^) showed similar pRB protein levels to WT hiPSCs (Fig. S3B). The second one was established by reprogramming somatic cells of a patient with *RB1* heterozygous mutation (c.658C>G, p.Leu220Val) (Fig. S12A, B)^29^. Sanger sequencing confirmed that the same mutation present in hiPSCs (Fig. S12C). To analyze the pathogenic character of this mutation, mRNA splicing analysis of patient-specific hiPSCs was performed using cDNA reverse-transcribed from RNA (Fig. S12D). PCR products revealed that two types of alternatively spliced mRNA were generated in patient-specific hiPSCs, which a part of exon 7 or the entire exon 8 were skipped (Fig. S12E, F). Because of it, a decreased pRB protein level was observed (Fig. S12G, H).

When induced into ROs, the higher ratio of Ki67^+^ cells in ATOH7^+^ cells was identified in patient-specific ROs at day 50 (Fig. 6A, B), and more ATOH7^+^ cells were identified as well (Fig. 6C). However, the low pRB level did not alter the proliferation of CPs, RGCs and RPCs (Fig. 6A, S13A). After 90 days, patient-specific ROs were significantly larger than WT and *RB1*^W/Puro^ ROs (Fig. 6D, E). In addition, the retinal lesion was observed in high frequency in patient-specific ROs with no proliferative CPs (Fig. 6D-G), which suggested that the low pRB level would be more likely to induce retinal lesions, which might mimic the cellular features of retinocytoma^30^. In the normal part of retina, RPCs were observed in both ROs at day 90 (Fig. S13B). Importantly, more photoreceptors and a higher cone-rod ratio were identified in patient-derived ROs (Fig. 6 H-J). The normal retina eventually developed into mature stage consisting of matured photoreceptors and other retinal cell types (Fig. 6K, L, S13C). Collectively, these results confirmed that although the monoallelic loss of *RB1* is insufficient for CPs proliferation, the enrichment of ATOH7^+^ cells in *RB1*^+/−^ ROs induces the retinal lesion by generating more non-proliferative cones.

**Fig. 6.**
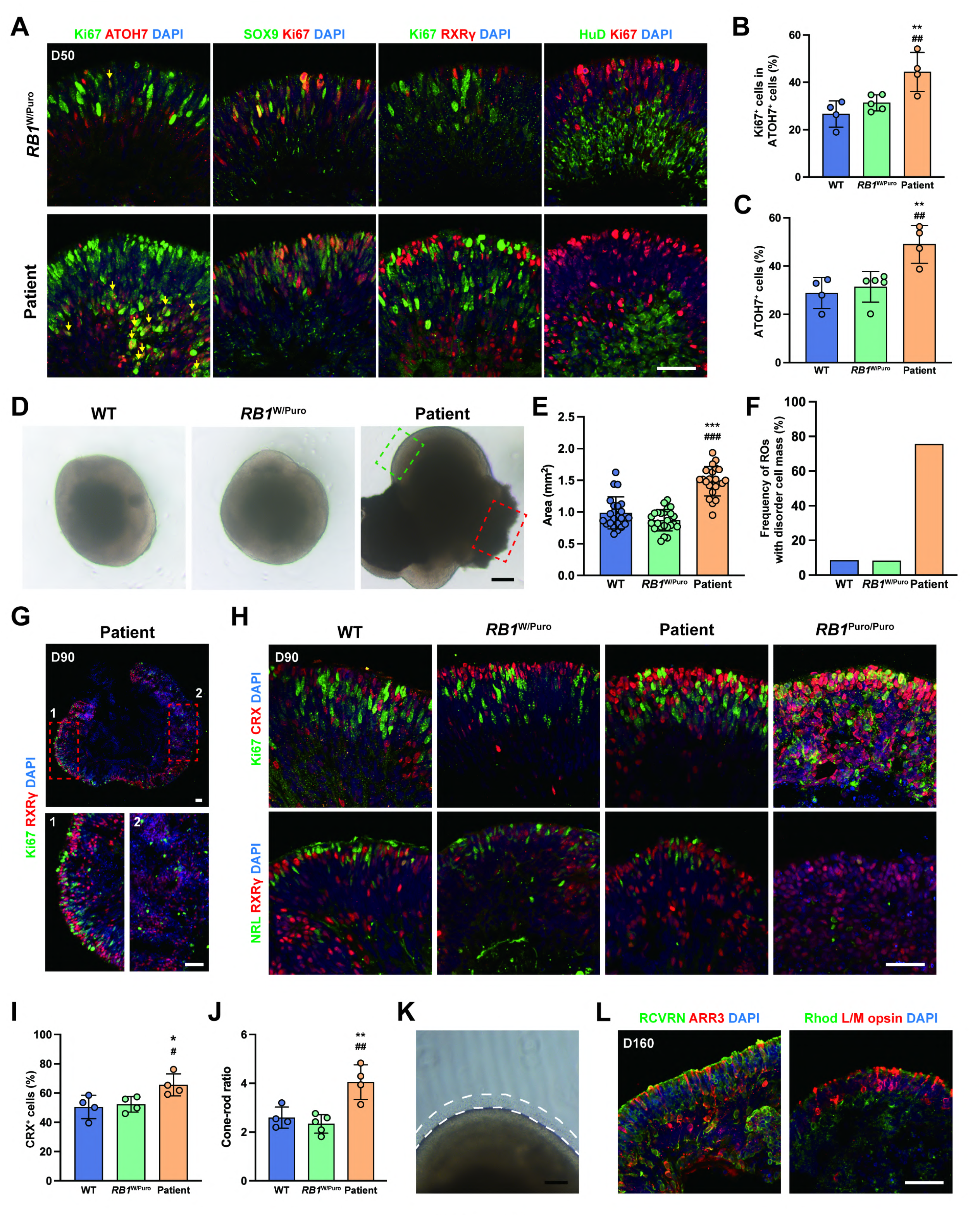
pRB protein level-dependent developmental defects in *RB1*^+/−^ Ros. (A) Representative immunostaining for SOX9, RXRγ, HuD and Ki67 in *RB1*^W/Puro^ and patient-specific ROs at day 50. Arrows indicate the Ki67^+^ cells in ATOH7^+^ cells. (B) Quantification of Ki67^+^ cells in ATOH7^+^ cells in different types of ROs at day 50. Data represent mean ± SD. ** P < 0.01 vs. WT; ## P < 0.01 vs. *RB1*^W/Puro^, n = 4-5. (C) Quantification of the ratio of ATOH7^+^ cells in different types of ROs at day 50. ** P < 0.01 vs. WT; ## P < 0.01 vs. *RB1*^W/Puro^, n = 4-5. (D) Representative microscopic images of WT, *RB1*^W/Puro^ and patient-specific ROs at day 90. Dashed green box indicate the neural retina. Dashed red box indicate the retinal lesion in patient-specific ROs. (E) Quantification of size of different types of ROs at day 90. Data represent mean ± SD. *** P < 0.001 vs. WT; ### P < 0.001 vs. *RB1*^W/Puro^, n ≥ 20. (F) Quantification of frequency of ROs with retinal lesion in different types of ROs at day 90, n ≥ 20. (G) Representative immunostaining for RXRγ and Ki67 in patient-specific ROs at day 90. Region 1 indicates the normal neural retina. Region 2 indicates the disorder cell mass. (H) Representative immunostaining for CRX, Ki67, RXRγ and NRL in WT, *RB1*^W/Puro^, patient-specific and *RB1*^Puro/Puro^ ROs at day 90. (I) Quantification of the ratio of CRX^+^ cells in different types of ROs at day 90. Data represent mean ± SD.* P < 0.05 vs. WT; # P < 0.05 vs. *RB1*^W/Puro^, n = 4-5. (J) Quantification of the ratio of cones/rods in different types of ROs at day 90. Data represent mean ± SD. ** P < 0.01 vs. WT; ## P < 0.01 vs. *RB1*^W/Puro^, n = 4-5. (K) Representative microscopic images of patient-specific ROs at day 160. (L) Representative immunostaining for RCVRN, ARR3, Rhodopsin and L/M opsin in patient-specific ROs at day 160. Scale bars = 50 μm (A, G, H, L), 200 μm (D) and 100 μm (K).

### 7. Multi-omics analysis of *RB1*^−/−^ ROs identifies a potential therapeutic target

We explored new potential therapeutic targets using multiple sequence datasets acquired from the RO-derived Rb models. ATAC-seq was performed to assess the differences in chromatin accessibility^31^. *RB1*^−/−^ ROs showed more chromatin-accessible regions than WT ROs (Fig. 7A, S14A), with the majority of the open regions being located in distal intergenic regions and introns, which were enriched in pathways related to cancer and cell cycle (Fig. 7B, C). Conversely, a smaller proportion of distal intergenic regions were open in WT ROs (Fig. S14B).

**Fig. 7.**
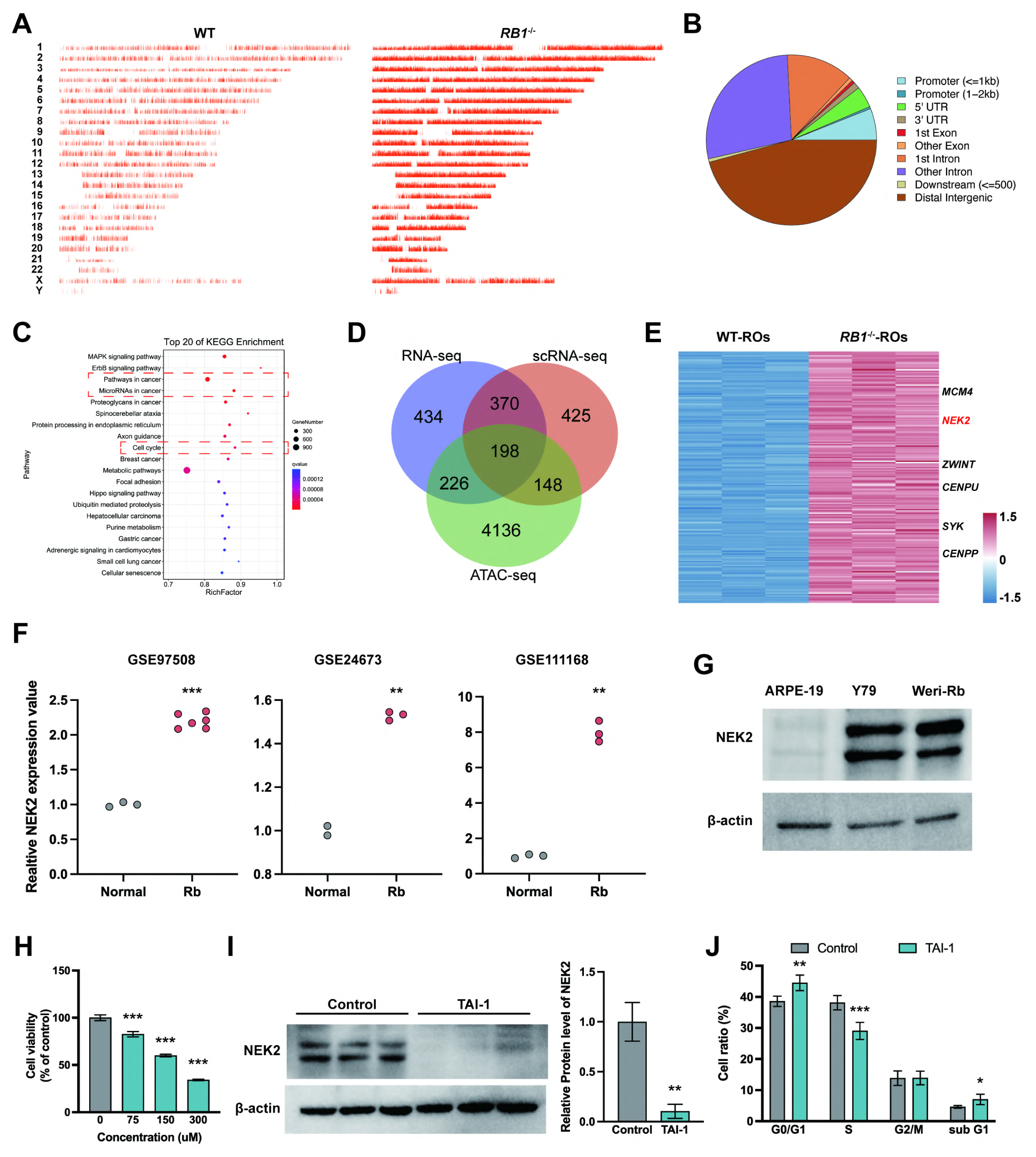
Exploration of potential therapeutic target for Rb from multi-omics analysis of *RB1*^−/−^ Ros. (A) Visualization of peaks over chromosomes in WT and *RB1*^−/−^ Ros (B) Distribution of open chromatin across the whole genome in *RB1*^−/−^ Ros (C) The top ten pathways enriched among the genes with open chromatin by KEGG pathway analysis. (D) Venn diagram showing the collective up-regulated genes by the integrative analysis of bulk RNA-seq, scRNA-seq and ATAC-seq. (E) Heatmap of the collective up-regulated genes from three data set in WT and *RB1*^−/−^ ROs. (F) Scatterplot plot of *NEK2* expression among retinoblastoma and normal retina sample in GSE97508, GSE24673 and GSE111168. *** P < 0.001, ** P < 0.01 vs. normal retina, n = 2-6. (G) Western blot analysis of NEK2 in Rb cell line, Y79 and Weri-Rb-1 as well as normal retinal cell ARPE-19. (H) Cell viability of Y79 cells after 72 hrs treatment with TAI-1 in different concentration by CCK8. Data represent mean ± SD. *** P < 0.001 vs. control, n = 6. (I) Western blot analysis of NEK2 after TAI-1 treatment in Y79 cells. Data represent mean ± SD. ** P < 0.01 vs. control, n = 3. (J) Cell cycle distribution of Y79 cells with TAI-1 treatment. Data represent mean ± SD. * P < 0.05, ** P < 0.01, *** P < 0.001 vs. control, n = 6.

Next, three datasets (RNA-seq, scRNA-seq, and ATAC-seq) were analyzed to explore potential therapeutic targets. DEGs between WT and *RB1*^−/−^ ROs were identified on day 80 (Fig. S14C). From the scRNA-seq data, genes with higher expression in Rb cells (Rb and P-Rb) were selected. After merging, 198 common upregulated genes were found (Fig. 7D, E). Among these, 10 hub genes were identified, including *NEK2* (Fig. 7E, S14D, E). This gene encodes a kinase that is involved in mitotic regulation^32^. Clinical Rb samples and human Rb cell line showed higher expression than normal retina (Fig. 7F, G). TAI-1, a small molecule that potentially induced NEK2 degradation, was used to explore the therapeutic effect for Rb *in vitro*^33^. TAI-1 showed no obvious toxicity in ARPE-19 cells at concentrations from 75 to 300 nM (Fig. S14F), but reduced the viability of Rb cell lines (Fig. 7H, S14G) and inhibited the protein level of NEK2 (Fig. 7I, S14H). Cell cycle analysis revealed that TAI-1 treatment induced S phase arrest and apoptosis in Rb cells (Fig. 7J, S14I). Overall, these results reveal that the RO-derived Rb models are helpful in identifying key genes involved in Rb development.

## Discussion

In this study, we generated *RB1* inactivated ROs in different genetic backgrounds to explore the role of pRB in Rb tumorigenesis and retinogenesis. It is the first time that an earlier type of CPs was identified as another cell-of-origin. In addition, we found a potential therapeutic target for Rb using multi-omics analysis. These novel findings provide new insights into the molecular and cellular mechanisms of Rb tumorigenesis.

Xenograft model is necessary to confirm the tumorigenesis of RO-derived cells^34,35^. Although several ROs-based Rb models have been developed, only Dyer’s group and Jin’s group explored the tumorigenicity *in vivo*^18,19^. Our data that using *RB1*-inactivated ROs from day 70 also confirm the tumorigenicity. However, Norrie et al. reported that tumors were identified over 1 year after xenograft, whereas the latency time from Liu et al. and our results were about 3 months. Because most of the cells in ROs at 40 days are RPCs, which may be not the cell type directly affected by *RB1* loss. Thus, an in-dish model closely mimicking human retinal development is a better model to explore when or how Rb initiates. Moreover, it is the first time that we observe the malignant transformation with consecutive xenograft of ROs-derived cells, which highlight the important role of the microenvironment in tumor progression^36,37^.

Despite considerable progress in uncovering the function of pRB in tumorigenesis, it is still unclear if pRB also plays the important role in normal development of human retina. In mouse model, although embryos with homozygous mutation of *RB1* die between day 14 and 15 of gestation, the development of retina seems to be normal in early retinogenesis^13,38^. Here, we confirmed that pRB loss did not make RPCs lose the property to develop into CPs and RGCs, but led to the developmental defect of late retinogenesis. Because pRB is necessary for RPCs to keep its cell cycle normally^6^, one potential reason is that *RB1* inactivation may directly alter early RPCs to re-enter cell cycle to generate late RPCs. Additionally, an indirect way to influence proliferation of RPCs by the loss (RGCs) or gain of specific cells (nRPCs) is another possible reason^39^. Because we cannot identified a obvious defect of RPCs in *RB1*^+/−^ ROs, we consider that the homozygous mutation of *RB1* gene would be a key factor that affect the development of late RPCs.

Although several reports imply that ARR3^+^ maturing CPs is the cell-of-origin of Rb, the *RB1* loss also contributes to early CPs proliferation^21^, which arises the possibility that Rb may initiates in an earlier stage of retinogenesis. In this study, we identified that the ATOH7^+^ nascent CP is an early stage of retinal cell in ROs that was sensitive to *RB1* loss. scRNA-seq data reveals this specific cell type may directly develop into a proliferative type of Rb. According to the results that *RB1*-null ATOH7^+^ RGCs also gain proliferative capacity in early retinogenesis^40^, some genes up-regulated in nRPC-stage may be another factor that activate the proliferation potential after pRB loss in CPs. However, because such cells only can be identified in *RB1*-null ROs before day 70, it is difficult to identify them in clinical samples. With this observation, we consider that the tumor initiation should be correlated to the time and the cell types that “second hit” occurred. If the “second hit” is before the emergence of ATOH7^+^ CPs, the tumorigenesis can initiate in the early retinogenesis^32^. Otherwise, if the “second hit” or “both hit” occurred in later stage, the tumor initiation would be from the maturing CPs.

To date, little is known about the positive relationship between germline mutations in *RB1* gene and the likelihood of developing Rb or retinocytoma^41^. One potential reason is that when the first hit occurred, the expression of normal pRB was reduced or pRB was inactivated^42^. In the ROs with low expression of pRB, the higher ratio of Ki67 was identified only in ATOH7^+^ cells but not in RXRγ^+^ or HuD^+^ cells, which confirmed that nRPCs were the more sensitive cell type to pRB. The proliferation of nRPCs resulted in more non-proliferative CPs and lead to higher frequency of retinal lesion, which was considered as a phenotype of retinocytoma. These results not only identify a retinal cell type that more sensitive to pRB than CPs, but also explore the mechanism that how germline mutations of RB1 develop retinocytoma for the first time. Developing new therapies saving vision and eyeballs for patients with Rb is urgently needed^43^. In recent years, many potential therapeutic targets have been explored in patient-derived tumors^34,44–47^. We constructed a multi-omics dataset to identify a potential therapeutic target of Rb. As a member of the never-in-mitosis A (NIMA) protein, NEK2 is a conserved centrosome kinase that plays crucial roles in cell cycle progression and differentiation^32^. The abnormal expression of NEK2 has been identified in a wide variety of human cancers and enhances malignancy via several signaling pathways^48–50^. From online data sources, we also identified the overexpression of *NEK2* in Rb tumors from patients and confirmed it using Rb cell lines. In addition, we found that TAI-1 is a potential therapeutic agent for treating Rb. Our results provide a druggable target not only for Rb but also for other cancers with NEK2 overexpression.

In conclusion, we successfully develop Rb model using *RB1*-knockout ROs. Completed *RB1* loss initially induces the proliferation of ATOH7^+^ nRPCs in early retinogenesis, which contributes to generate more CPs and RGCs and leads to development defect of late retinogenesis in pRB-null ROs. The death-resistant CPs form tumor tissue with mature cone marker expression. Additionally, the abnormal proliferation of ATOH7^+^ nRPCs also identified in *RB1*^+/−^ ROs with pRB down-regulation, which may be the initiation of retinocytoma. With the multi-omics analysis of pRB-null ROs, a potential therapeutic target is identified. Our study not only contribute to identifying a new cell-of-origin of Rb, but also facilitating the development of new efficacious therapies.

## Materials and methods

### Clinical samples from Rb patients

Human retinal tissue samples were obtained from Zhongshan Ophthalmic Center, Sun Yat-sen University with patient informed consent and approved by Ethics Committee. Sections for immunohistochemistry (2021000019, 2021000020, 2021000028) and for Hematoxylin/Eosin staining (2020111315) were obtained from primary tumor tissues.

### Maintenance of cell lines

All cells were maintained at 5% CO_2_ in a 37°C incubator. Y79 and Weri-Rb-1 retinoblastoma cell lines were cultured with RPMI Medium 1640 (Gibco, #C11875500BT) with 20% fetal bovine serum (FBS) (Gibco) and 1% penicillin-streptomycin (Gibco, #15140-122). ARPE19 cell line was cultured with DMEM/F12 (Gibco, #C11330500BT) with 10% FBS and 1% penicillin-streptomycin. The medium was changed every second day.

### Maintenance of hiPSCs

BC1 hiPSC was a gift from Professor Linzhao Chen of University of Science and Technology of China^51^. Gibco hiPSC was purchased from Thermo Fisher Scientific. hiPSCs were cultured according to the previous method^15,25^. Patient-derived hiPSC was generated by reprograming of somatic cells from a retinocytoma patient. Cells were grown on 6 well plates coated with Matrigel (Corning, #354277) in mTeSR1 medium (Stem Cell Technologies, #85850) at 5% CO_2_ in a 37°C incubator. Medium was refreshed every second day. When it grew to 80% confluency, hiPSCs were treated with 0.5 mM EDTA (Invitrogen) for passage.

### Retinal organoids differentiation

Retinal differentiation was performed according to previous protocols^15^. Briefly, hiPSCs were digested into small clumps and cultured in suspension with mTeSR1 supplemented with 10 μM Blebbistatin (Sigma-Aldrich) to form embryoid bodies (EBs) on day 0. Then culture medium was switched into 3:1 ratio of mTeSR1/Neural induction medium (NIM) which containing DMEM/F12, 1% N2 supplement (Invitrogen), 1% non-essential amino acids (NEAA) (Gibco), 2 μg/mL heparin (Sigma-Aldrich) on day 1, 1:1 ratio of mTeSR1/NIM on day 2 and 100% if NIM on day 3. EBs were plated onto Matrigel-coated dished between day 5 to day 7. On day 16, the culture medium was changed to retinal differentiation medium (RDM) containing 72% of DMEM Basic (Gibco, #C11995500BT), 24% of DMEM/F12, 2% B27 supplement without vitamin A (Gibco), 1% NEAA and 1% antibiotic-antimycotic (Gibco). From week 4 to week 6, Neuron retina domains in good condition were manually detached with a sharpened Tungsten needle and transferred to suspension culture for further induction of retinal organoids. For long time culture, organoids were cultured with retinal culture medium (RCM) containing RDM, 10% FBS, 100 uM Taurine (Sigma) and 2 mM GlutaMax (Gibco) within 1 week after detachment. From week 13 onwards, the medium was switched to RCM2, which B27 supplement was replaced by N2 supplement in RCM. Medium was changed every 3 days.

### Xenografts and in vivo imaging

The experimental procedures were approved by the ethics committee of Sun Yat-sen University. For the first round of transplantation, ROs between day 60 to day 70 were dissociated into single cells as previous report using papain dissociation system (Worthington Biochemical)^25^, and the cells were temporarily centrifuged and resuspended in DMEM/Basic medium. Prior to cell injection, 4-6 weeks old NOD/SCID mice were anaesthetized with intraperitoneal sodium pentobarbital injection (50 mg/kg, Sigma), followed by anaesthesia with 2.5% phenylephrine hydrochloride solution and pupil dilation with 5 mM local anaesthetic proparacaine hydrochloride. The cells were then injected into the subretinal space or vitreous cavity with 1.5 µL suspension (containing 1.0-2×10^5^ cells) per eye using a 2.5 µL microinjector and 35G syringe needle (Hamilton, Switzerland). Tobramycin was used to prevent ocular infection. Spectral domain optical coherence tomography (SD-OCT) with an optical coherence tomography scanner (Heidelberg Engineering, SpectralisOCT, Germany) and anterior chamber photography with a slit lamp digital imaging system (Topcon, SL-D7/DC-3/IMAGEnet, Japan) were used to track and monitor the grafts. After 13 weeks of injection, the mice were executed and tumor-bearing eyes were used for further analysis. In the second round of xenograft, the tumor cells that from first xenograft were dissociated into single cells with 0.5 mM EDTA for 5 min and then 1-2×10^5^ cells for each eye were used for transplantation. The procedure for the second round of xenograft followed that of the first, with tumor cells collected and analyzed after 13-15 weeks.

### Teratoma formation and analysis

The experimental procedures were approved by Sun Yat Sen University’s Ethics Committee. Experiment was performed as previous reports. 1-1.5×10^6^ hiPSCs with 30% Matrigel were injected into the hind limb of NOD/SCID mice. After 6–8 weeks, teratomas were dissected from animals, fixed in formalin, embedded in paraffin, sectioned and stained with Hematoxylin and Eosin (H&E). Images were taken using a Zeiss Axio Scan. Z1 Slide Scanner (Carl Zeiss, Jena, Germany).

### Targeting, donor plasmid construction

Two *RB1*^−/−^ hiPSC lines were generated by CRISPR/Cas9 gene editing, with human BC1 and Gibco hiPSC as parents. For the BC1 knockout cell line, a single guide RNA (sgRNA) was designed by the online website: http://crispor.tefor.net/ (the sgRNA sequence is GGTGGCGGCCGTTTTTCGGG), which targeting *RB1* exon 1 CDS (Coding Sequence) region was inserted into pSpCas9(BB)-2A-Puro (PX459 V2.0, Addgene) vector to generate the targeting plasmid PX459-RB1-sgRNA. Similarly, the same sgRNA was inserted into pSpCas9(BB)-2A-GFP (PX458, Addgene) vector to establish the targeting plasmid PX458-RB1-sgRNA. hPGK-Puro-polyA resistance cassette was flanked by left and right homologous arms. The above fragment was then inserted into pMD19-T (Takara, #D102A) vector to generate the donor plasmid pMD19-LA-Puro-RA.

### Cell transfection and resistance selection

Electroporation was carried out as previously described^25^. In brief, hiPSCs were dissociated into single cells using Accutase (Gibco, #A1110501) at 37°C for 5 min, Approximately 15 µg PX459-RB1-sgRNA plasmid for BC1 hiPSC or 10 µg PX458-RB1-sgRNA and 10 µg pMD19-LA-Puro-RA plasmid for Gibco hiPSC were used for further nucleofection. After 3 days of transfection, resistant single cell clones were selected with puromycin (Solarbio) at a concentration of 0.4 µg/mL for BC1-*RB1*^−/−^ hiPSCs or 1 µg/mL for Gibco-*RB1*^Puro/Puro^ hiPSCs. The above clones were amplified and used for further validation.

### Mutation analysis

Genomic DNA was extracted using an Animal Genomic DNA Rapid Extraction Kit (Beyotime). PCR was performed using primers (primer VF1, VR1 for BC1-*RB1*^−/−^ hiPSC, VF2, VR2, VF3, VR3 for Gibco-*RB1*^Puro/Puro^ hiPSCs and Gibco-*RB1*^w/Puro^ hiPSCs, see sequence details in Table S3) based on which exon was mutated in the sample. Mutation was verified by Sanger sequencing.

### RT-PCR analysis

Total RNA was extracted from hiPSCs using Trizol Reagent (Sigma). Then, 1 μg of total RNA was reverse-transcribed using the PrimeScript™ RT reagent Kit (Takara, #RR047A) to synthesize cDNA. The VF4/VR4 primer pair was used to perform PCR to verify the mutant effect (see sequence details in Table S3). The resulting PCR products were used for further Sanger sequencing.

### Karyotype analysis

G-band staining of chromosomes was used for karyotype analysis. hiPSCs were treated with 20 µg/mL colchicine for 1 h. Cells were then digested with 0.5 mM EDTA for 5 min, resuspended and incubated in 0.075 M KCl solution for 20 min, and then fixed with a 3:1 methanol/acetic acid mixture for 10 min. After dropping onto cold slides for 2 h, chromosomes were observed under an OLYMPUS BX43 Microscope and analyzed with Applied Spectral Imaging system.

### Cell cycle analysis

Cell cycle profiles were analyzed by the Cell Cycle Staining Kit (MultiScience, Hangzhou, China) according to the manufacturer’s instructions. Briefly, cells were dissociated into single cells. After washing twice with PBS, cells were fixed with cold 75% ethanol overnight. The cells were then washed, treated with RNase and stained with PI in the dark for 30 min. The percentage of cells in the sub G1, G0/G1, S and G2/M phases of the cell cycle was analyzed using a BD LSR Fortessa cytometer (BD Biosciences).

### Immunocytochemistry

For cryosection analysis, samples were fixed in 4% paraformaldehyde (PFA) for 30 min and dehydrated in sucrose solution. After embedding in OCT compound (Sakura Finetek Japan, Tokyo, Japan), tissues were sectioned at a thickness of 12-16 μm. For flatmounts analysis, samples were fixed in 4% PFA for 15 min. Then samples were blocked and permeabilized in 0.4% TritonX-100 with 1% Donkey serum for 1 hour at room temperature and then incubated with primary antibody at 4°C overnight. The following day, samples were rinsed with PBS and subsequently incubated with secondary antibodies for 1 hour at room temperature in the dark. Nuclei were stained with DAPI solution (Dojindo). Images were obtained using the Zeiss LSM 880 confocal microscope (Carl Zeiss). Antibody information is detailed in Table S4. Quantification of positive cells was performed using ImageJ or QuPath software^55,56^. Quantification of retinal dyslamination was performed according to recent report^57^.

### Hematoxylin and Eosin staining

Tissue and ROs were fixed in formalin, embedded in paraffin, sectioned and stained with Hematoxylin (Biosharp, Hefei, China) and Eosin (Beyotime Biotechnology, Shanghai, China) (H&E). Images were taken using a Zeiss Axio Scan. Z1 (Carl Zeiss, Jena, Germany).

### EdU labelling

EdU labeling was performed using Click-iT EdU Imaging Kits (Invitrogen). ROs at the indicated time were cultured in RCM in the presence of 10 μM EdU for 24 h before collecting. After fixation, dehydration, embedding, and segmentation, EdU was detected by Click-iT assay according to the manufacturer’s protocol.

### CCK8 assay

CCK8 (Cell Counting Kit-8) colorimetric assays were performed to detect the cell viability in ARPE19, Y79 and Weri-Rb-1 cells according to the manufacturer’s method (GLPBIO). Briefly, approximately 1×10^4^ cells were seeded in 96-well plate and incubated with/without different concentration of TAI-1 (ApexBio). After 72 h, CCK8 reagent was applied to each well and incubated for 2 h at 37°C. The absorbance at 450 nm of each well was measured using a multimode microplate reader (Synergy H1; BioTek) to calculate the relative cell viability.

### Western blot

hiPSCs, Retinoblastoma cell lines, ARPE19 cell line or ROs were harvested and washed with 1×PBS, with the addition of cold RIPA Lysis Buffer (Beyotime, #P0013B) and 1×PMSF (Beyotime, #ST507), followed by incubation on ice for 30 min in a shaker. The lysate supernatant was retained after 15 min centrifugation (4 °C) for SDS-PAGE. The BCA Protein Assay Kit (Beyotime) was used to determine the protein concentration of each sample. Approximately 10 to 20 µg of proteins from cells or organoids lysates were mixed with Loading Buffer (Beyotime, #P0015L), incubated at 95-100 °C for 10 min and separated on 10% Bis Tris Gel in Tris-Glycine Running Buffer (Sangon, #C520001). The proteins were transferred to 45 μm pore size PVDF membranes (Millipore, #HVLP02500) by Apparatus (Bio-Rad). Membranes were blocked with 5% non-fat milk solution or 5% BSA for 1 h, then incubated with primary antibodies (see Table S2 for details) at 4 °C overnight, followed by incubation with secondary antibodies for 1 h at room temperature using horseradish peroxidase-conjugated anti-rabbit or anti-mouse IgG. The signal was generated using Immobilon Western Chemilum HRP Substrate (Millipore, #WBKLS0100) and visualized by Bio-Rad ChemiDoc Touch system or Tanon 5200.

### Transmission electron microscope (TEM)

For TEM analysis, ROs and tissues were dehydrated by passing through a graded series of ethanol solutions and embedded in epoxy resin. Samples were then cut into ultrathin sections with an ultra-microtome, stained with 1% uranyl acetate and lead citrate, and then imaged with an electron microscope (FEI Europe, Eindhoven, Netherlands).

### Bulk RNA-seq analysis

ROs or hiPSCs were collected for RNA extraction. Total RNA was isolated with Trizol reagent (Invitrogen) according to the manufacturer’s method. RNA quality was assessed using Agilent 2100 bioanalyzer (Agilent Technologies). mRNA was enriched using Oligo(dT) beads and then broken into short fragments. The fragments were further reverse transcribed into cDNA using NEBNext Ultra RNA Library Prep Kit for Illumina (New England Biolabs). The cDNA was further repaired at the ends and amplified by PCR to generate a sequencing library. RNA sequencing was performed by Illumina Novaseq6000 Sequencer.

Raw data was processed using the fastp tool (version 0.18.0)^52^. Reads containing poly-Ns, duplicate sequences, and low-quality sequences were removed to obtain high-quality clean reads. The remaining clean reads were further used in assembly and gene abundance calculations. Gene expression levels were quantified by RSEM 16 software. The fragment per kilobase transcript per million mapped reads (FPKM) value was calculated to quantify the expression levels of each gene. Correlation analysis was performed by R. correlation of two parallel experiments. Principal Component Analysis (PCA) was performed with R package g models (http://www.r-project.org/). Differential expression analysis was performed by DESeq2 R package (1.18.0) between two different groups and edgeR between two samples. Genes/transcripts with the false discovery rate (FDR) parameter below 0.05 and absolute fold change ≥ 2 were considered DEGs (different expression genes). Then, the DEGs were analyzed by Gene Ontology (GO) and Kyoto Encyclopedia of Genes and Genomes (KEGG) pathway enrichment. GO terms or pathways with a Q value ≤ 0.05 were defined as significantly enriched. Gene set enrichment analysis (GSEA) was further performed using GSEA and MSigDB software. Enrichment scores and p values were calculated in default parameters, and p < 0.05 was considered statistically significant. Data have been submitted to GEO database.

### ATAC-seq analysis

Nuclei suspensions were incubated in a Transposition Mix that includes a Transposase. The Transposase entered the nuclei and preferentially fragments the DNA in open regions of the chromatin. Simultaneously, adapter sequences were added to the ends of the DNA fragments. Incubated the transposition reaction at 37°C for 30 min. Immediately following transposition, the products were purified using a QIAGEN minielute kit, amplified and sequenced using Illumina NovaSeqTM 6000.

After filtering the clean reads, alignment reads by Bowtie 2^59^, peak scanning was performed by MACS. The read-enriched regions would be defined as a peak when log2 (fold enrichment above background) ≥ 3 &–log10 (p-value) ≥ 3. After that, the peak related genes were identified according to the genomic location information and gene annotation information of peak, peak related genes can be confirmed using ChIPseeker. Pathway enrichment analysis identified significantly enriched metabolic pathways or signal transduction pathways in peak related genes comparing with the whole genome background.

### Single-cell RNA-seq analysis

#### Library preparation and sequencing

Totally 15 - 20 ROs at day 80 were dissociated into single cells using papain enzyme according to the manufacturer’s instructions (Worthington Biochemical). Cell suspension was added to an equal volume of 0.4% trypan blue and the cells were counted using a Countess^®^ II Automated Cell Counter to adjust the concentration of live cells to the desired concentration (1000-2000 cells/μL). The cell suspension was then loaded onto a 10x Genomics GemCode Single-cell instrument to generate GEMs (Gel Beads-In-Emulsions). The libraries were prepared using Chromium Next GEM Single Cell 3’ Reagent Kits version 3.1 and sequenced in PE150 mode at an Illumina Novoseq6000 platform. A total of 9085 cells in WT ROs and 9520 cells in C5 *RB1*^−/−^ ROs were generated respectively.

### Cell clustering

CellRanger (version 3.1.0) software was used for quality control and expression quantification. Samples were de-multiplexed and aligned to the human reference genome GRCh38 to calculate cell-gene expression matrix.

The gene expression matrices of cells were submitted to Seurat (version 3.1.1) for further analysis^53,54^. Cells with unusually high numbers of UMIs (Unique Molecular Identifier) ≥8000 or mitochondrial gene percent ≥10% were filtered out. Cells with less than 500 or more than 6000 genes detected were also excluded. Additionally, doublet GEMs also filtered out by using DoubletFinder (v2.0.3)^55^. Seurat software’s LogNormalize method was used to normalize gene expression, FindVariableFeatures method was used to find intercellular high signature genes. Samples were then integrated using Canonical Correspondence Analysis (CCA) correction to remove batch effects. Finally, the data was normalized to Z-score using the Seurat’s ScaleData function, Cell clustering was performed by Louvain method with maximum modularity.

### Cell cycle analysis

The Seurat was used to assign the cell cycle score to each cell from 15 G1 phase genes, 20 S phase genes, 61 G2 phase genes and 64 M phase genes based on the previous reports^63^.

### Pseudo trajectory analysis

Single cell trajectory was analyzed using matrix of cells and gene expressions by Monocle 2 (Version 2.6.4)^56^. The high-dimensional cell expression data was projected into a two dimension space by a dimensionality reduction method (DDRTree) to order the cells (sigma = 0.001, lambda = NULL, param.gamma = 10, tol = 0.001), DEGs across different clusters (FDR<1e-5) were used to extrapolate the biological trajectory tree based on pseudo time.

### Online data sources and bioinformatic analysis

Three microarray or gene expression data set (GSE97508, GSE24673, GSE111168) were retrieved from Gene Expression Omnibus (GEO) database (https://www.ncbi.nlm.nih.gov/geo/). Totally 12 retinoblastoma samples and 8 normal retina samples were obtained from these three data sets.

The common DEGs from three data set (RNA-seq, scRNA-seq and ATAC-seq) were analyzed by the Search Tool for the Retrieval of Interacting Genes (STRING) database to investigate the interaction (https://www.string-db.org)^57^. To identify the hub genes in this network, the data from STRING was imported into the CytoHubba program (Cytoscape, https://cytoscape.org). The top 10 nodes with highest degrees were selected as hub genes.

### Statistical analysis

All experiments were repeated at least three times. Values were expressed as mean ± standard deviation (SD). Statistical analysis was performed with GraphPad Prism version 9.0. The statistical significance of difference was determined by unpaired t-test or one-way ANOVA followed by Dunnett’s test and p value below 0.05 was considered statistically significant.

## Acknowledgments

This work was supported partly from the National Natural Science Foundation of China (81970842 and 82172957); Science & Technology Project of Guangdong Province (2017B020230003); National Key Research and Development Program of the Ministry of Science and Technology (2017YFA0104101); a Joint grant from Science & Technology Project of Guangzhou and Zhongshan Ophthalmic Center, Sun Yat-sen University (202201020312); The Science & Technology Project of Guangzhou (202102010288) the Fundamental Research funds of the State Key Laboratory of Ophthalmology.

## Author contributions

K.Y., Y.W., P.X. and X.Z. designed the experiments. K.Y., Y.W., P.X., B.X., S.W., W.Z., G.G., X.S., S.Z., D.Z., X.S., S.Z., and F.G. performed the experiments. K.Y., Y.W., P.X., Y.Li, Y.Liu, J.W., R.S., and X.Z. analyzed and interpreted the data. X.Z. supervised the project. K.Y., Y.W., P.X. and X.Z. wrote the manuscript. All authors approved the finalized manuscript.

