## Supplemental information for "Longitudinal analysis of retinal cell state transitions in *RB1*-deficient retinal organoids reveals the nascent cone precursors are the earliest cell-origin of human retinoblastoma"

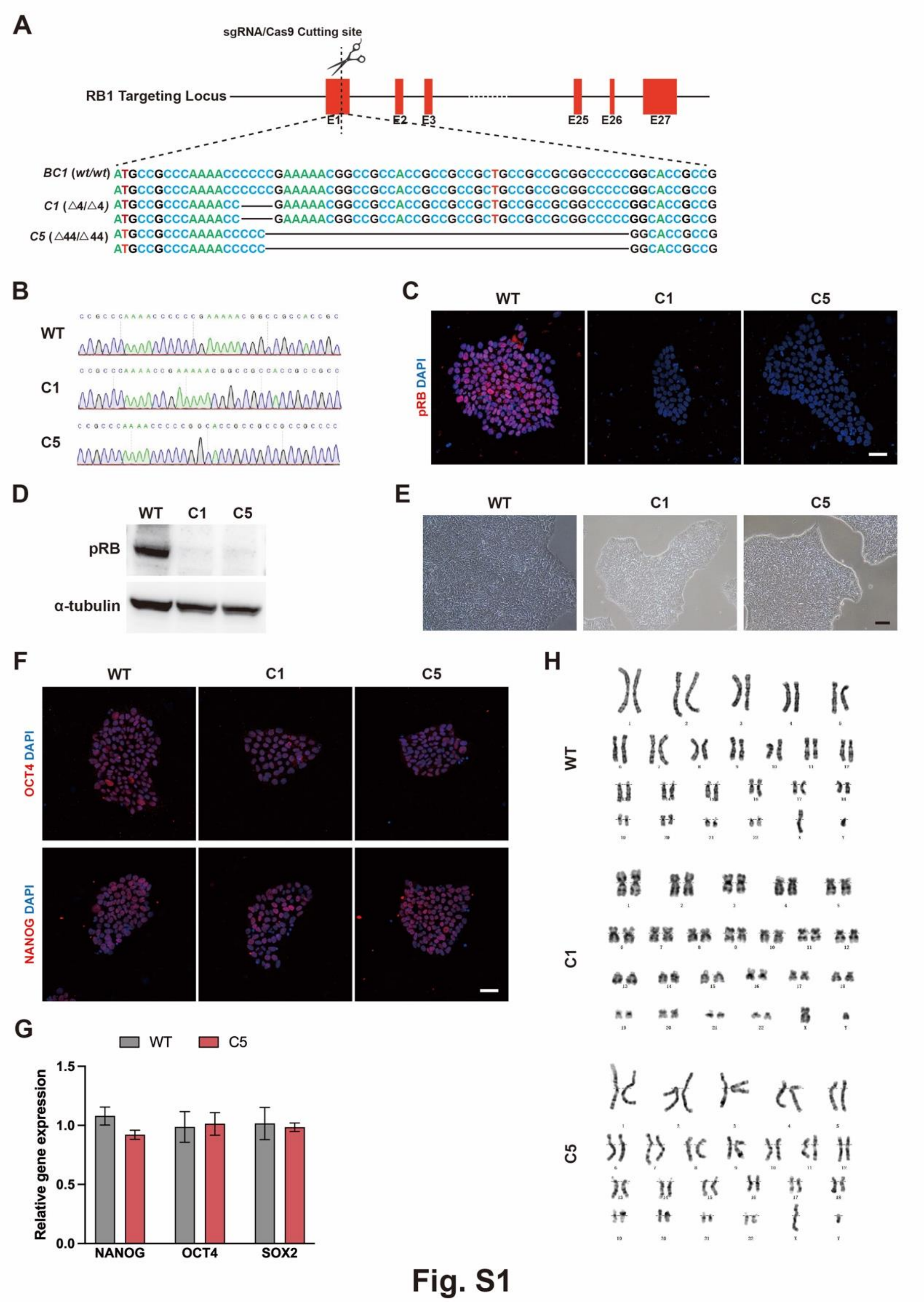


Fig. S1 Generation and characterization of *RB1*$^{-/-}$ hiPSCs

(A) Target strategy of generation of *RB1*$^{-/-}$ hiPSCs from BC1 hiPSCs.

(B) Sequences of targeted loci in different hiPSCs.

(C) Representative immunostaining images of pRB in WT and *RB1*$^{-/-}$ hiPSCs.

(D) Western blot analysis of pRB in WT and *RB1*$^{-/-}$ hiPSCs.

(E) Representative bright field images of WT and *RB1*$^{-/-}$ hiPSCs.

(F) Representative immunostaining images of OCT4 and NANOG in wild type and *RB1*$^{-/-}$ hiPSCs line.

(G) Representative expression of pluripotency genes in WT and *RB1*$^{-/-}$ hiPSCs, n = 3.

(H) Karyotype analysis of different hiPSC lines.

Scale bars = 200μm (C, E, F).

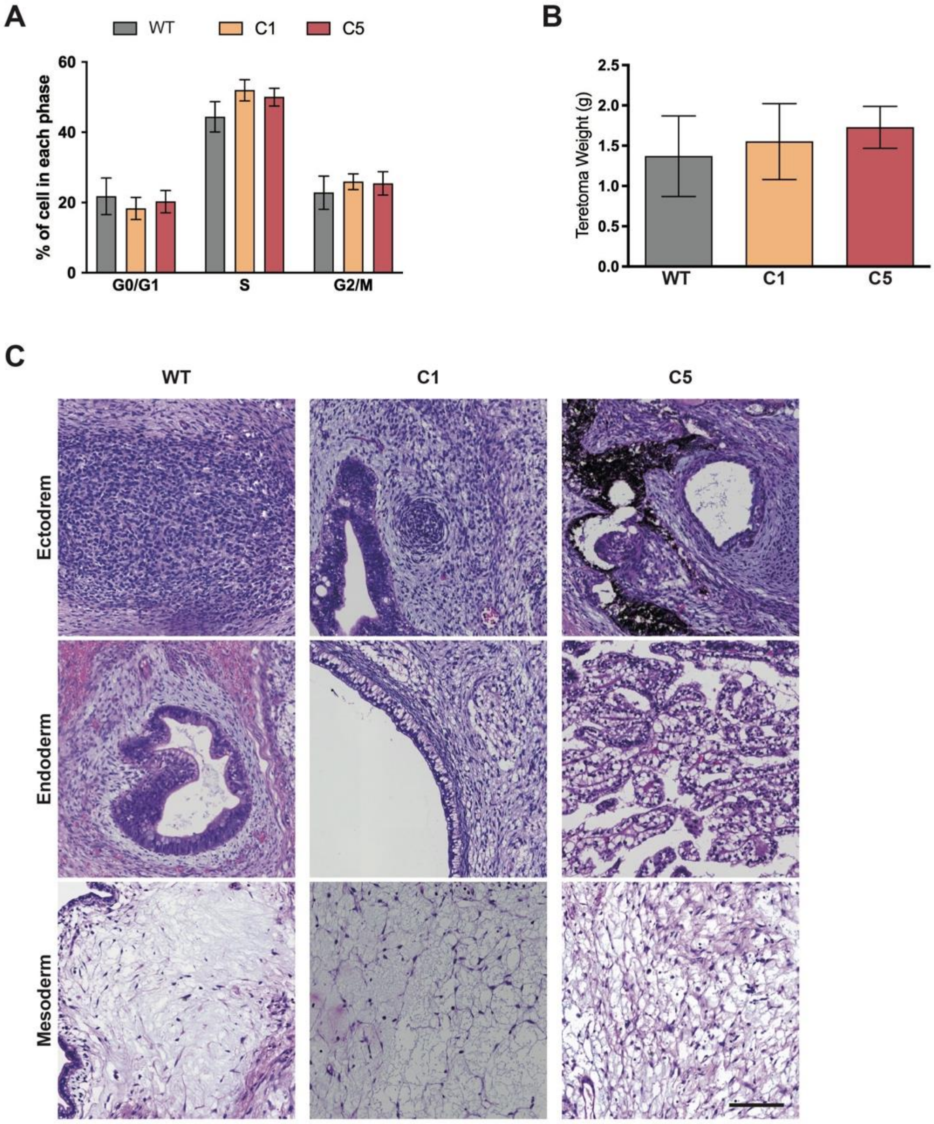


**Fig. S2**

<u>Fig.S2 Cell cycle and differentiation potential of *RB1*$^{-/-}$ hiPSCs.</u>

(A) Cell cycle distribution of WT and *RB1*$^{-/-}$ hiPSCs, n = 5-6.

(B) Quantification of weight of teratomas generated from WT and *RB1*$^{-/-}$ hiPSCs, n ≥ 5.

(C) Representative H/E staining images of ectoderm, endoderm and mesoderm-derived cells in teratomas generated from WT and *RB1*$^{-/-}$ hiPSCs.

Scale bars = 100 μm (C).

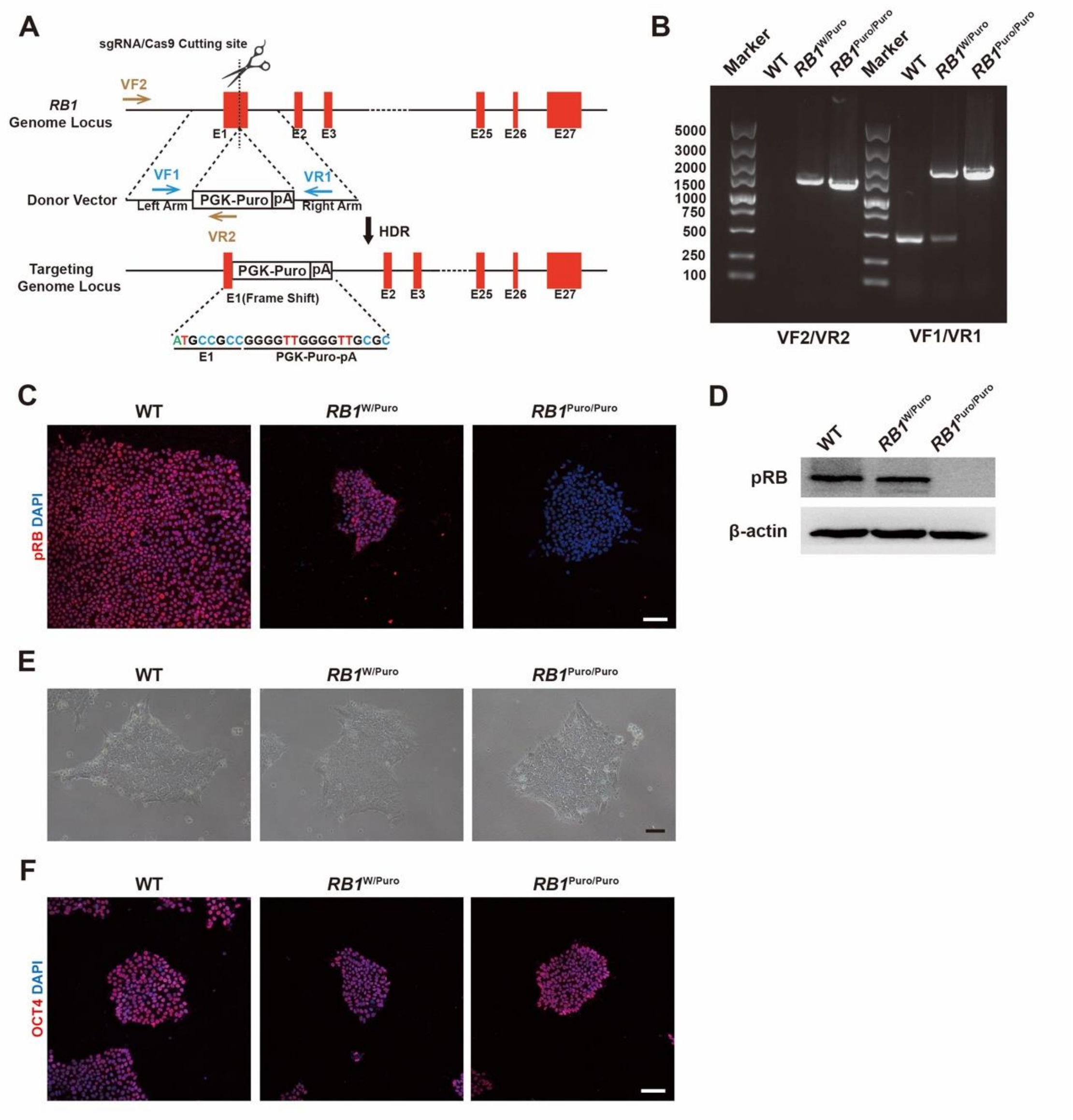


**Fig. S3**

Fig. S3 Generation and characterization of Gibco line-derived $RB1^{-/-}$ and $RB1^{+/-}$ hiPSCs

(A) Target strategy of generation of $RB1^{W/Puro}$ and $RB1^{Puro/Puro}$ hiPSC line from Gibco hiPSCs. Two primer pairs were designed to confirm the knockout of *RB1* gene and insertion of PGK-puromycin cassette.

(B) PCR products from wild type, $RB1^{W/Puro}$ and $RB1^{Puro/Puro}$ hiPSCs using VF1/VR1 primer pair (right) or VF2/VR2 primer pair (left). $RB1^{W/Puro}$ and $RB1^{Puro/Puro}$ hiPSC line have a longer product than WT hiPSC line using VF1/VR1 primer pair because of the insertion of PGK-puromycin cassette. $RB1^{W/Puro}$ hiPSC line has one more normal product. The PCR products were identified in $RB1^{W/Puro}$ and $RB1^{Puro/Puro}$ hiPSC line using VF2/VR2 primer pair, which confirmed the exist of PGK-puromycin cassette.

(C) Representative immunostaining for pRB in WT, *RB1*$^{W/Puro}$ and *RB1*$^{Puro/Puro}$ hiPSCs.

(D) Western blot analysis of pRB in WT, *RB1*$^{W/Puro}$ and *RB1*$^{Puro/Puro}$ hiPSCs.

(E) Representative bright field images of WT, *RB1*$^{W/Puro}$ and *RB1*$^{Puro/Puro}$ hiPSCs.

(F) Representative immunostaining images for OCT4 in WT, *RB1*$^{W/Puro}$ and *RB1*$^{Puro/Puro}$ hiPSCs.

Scale bars = 100 µm (C, F) and 200 µm (E).

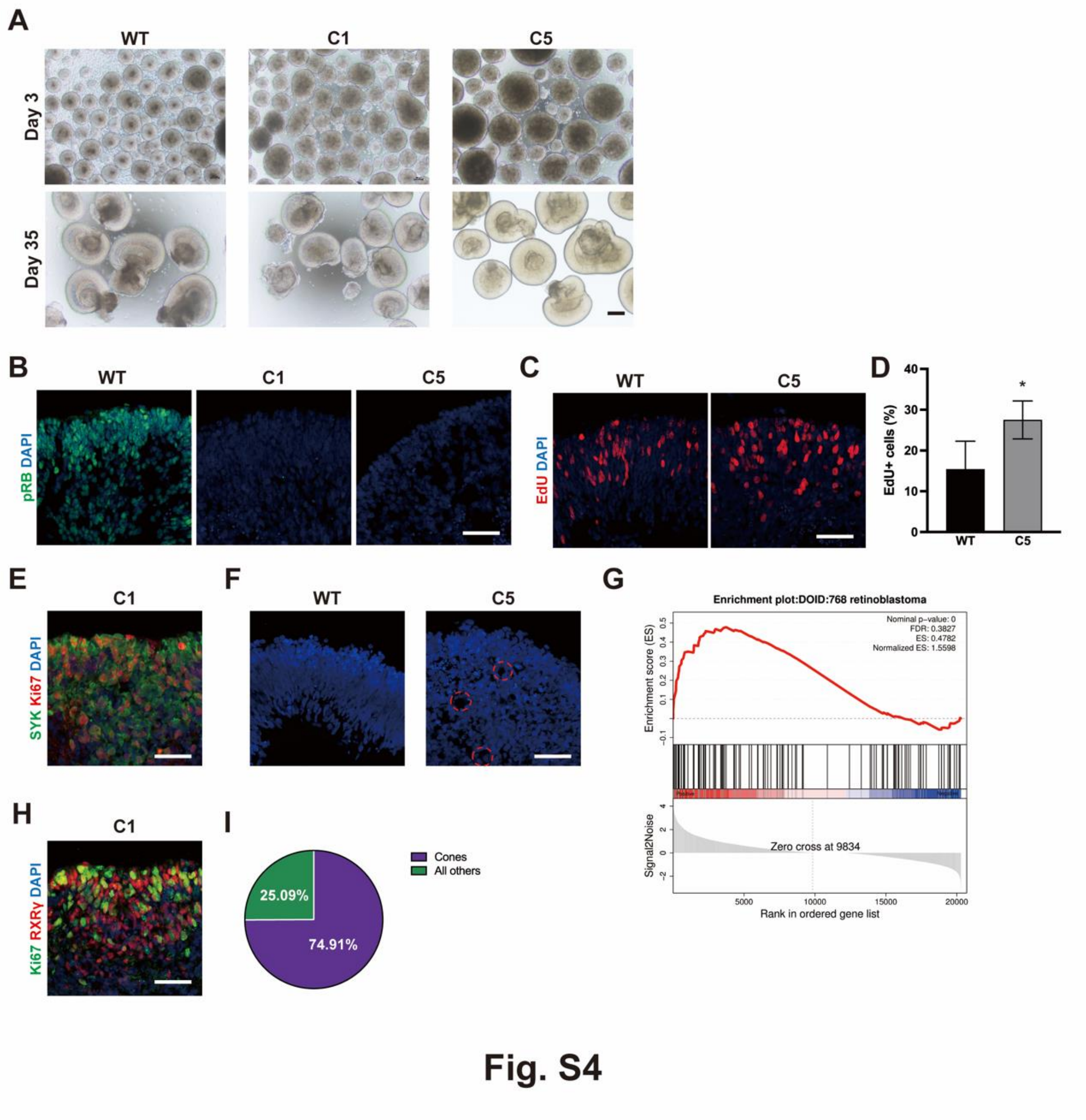


Fig. S4 Induction of ROs from *RB1*$^{-/-}$ hiPSCs

(A) Representative bright field images of EBs and early ROs in WT and *RB1*$^{-/-}$ ROs.

(B) Representative immunostaining images of pRB in WT and *RB1*$^{-/-}$ ROs at day 50.

(C) Representative EdU-labeling images in WT and *RB1*$^{-/-}$ ROs at day 70.

(D) Quantification of EdU-labeled cells at day 70. Data represent mean ± SD. * $P < 0.05$ vs. WT, n = 4.

(E) Representative immunostaining iamges of SYK and Ki67 in C1 *RB1*$^{-/-}$ ROs at day 90.

(F) Representative DAPI staining images in WT and *RB1*$^{-/-}$ ROs at day 90. The dash lines indicate the rosette structure.

(G) Up-regulation of Rb-related genes in *RB1*$^{-/-}$ ROs by Gene set enrichment analysis

(GSEA).

(H) Representative immunostaining for Ki67 and RXRγ in C1 *RB1*$^{-/-}$ ROs at day 90.

(I) Quantification of RXRγ$^{+}$ cells in C1 *RB1*$^{-/-}$ ROs.

Scale bars = 200 μm (A) and 50 μm (B, C, E, F, H).

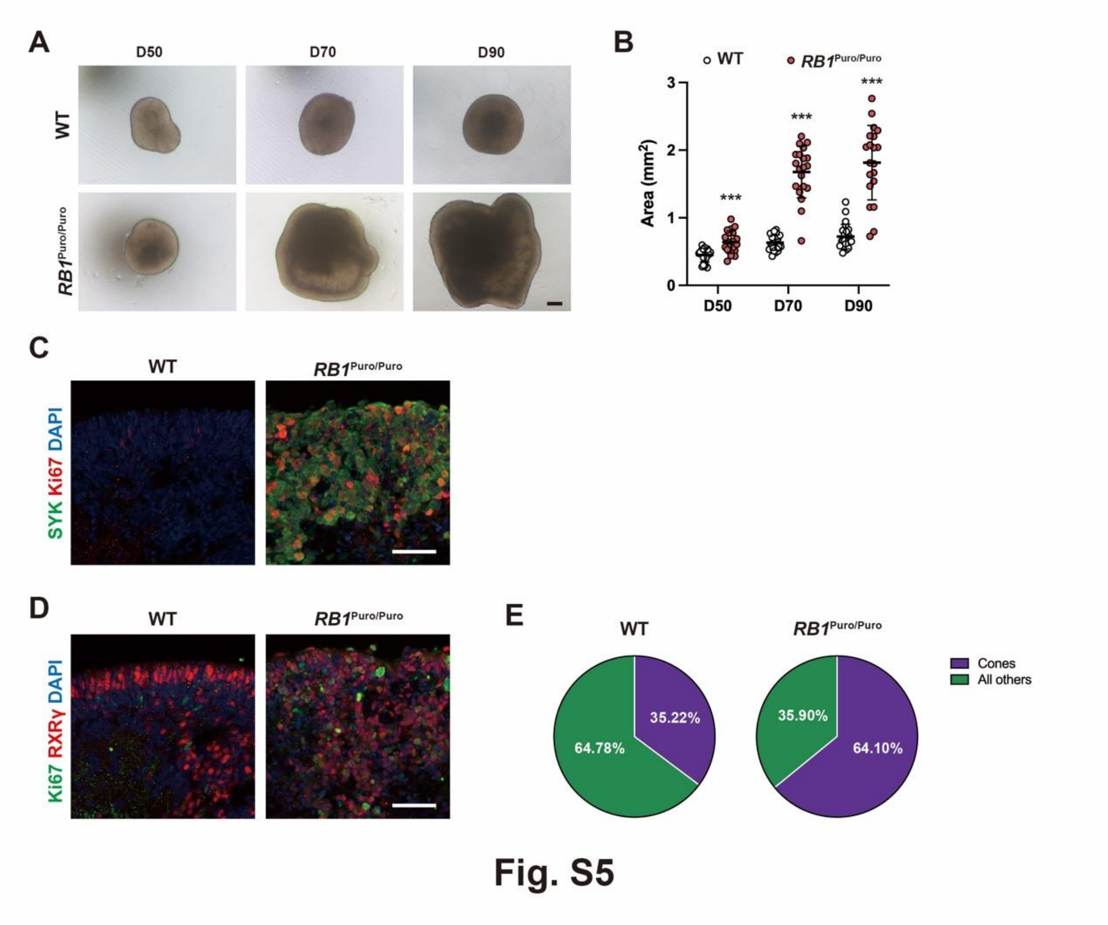


<u>Fig. S5 Reproducibility of tumorigenesis in *RB1*$^{Puro/Puro}$ ROs from Gibco hiPSC line</u>

(A) Representative microscopic images of WT and *RB1*$^{Puro/Puro}$ ROs from day 50 to day 90.

(B) Quantification of size of ROs in different culture phase. Data represent mean ± SD (n = 20-24). *** $P < 0.001$ vs. WT, $n \geq 20$.

(C) Representative immunostaining for SYK and Ki67 in WT and *RB1*$^{Puro/Puro}$ ROs at day 90.

(D) Representative immunostaining for Ki67 and RXRγ in WT and *RB1*$^{Puro/Puro}$ ROs at day 90.

(E) Quantification of RXRγ$^{+}$ cells in WT and *RB1*$^{Puro/Puro}$ ROs.

Scale bars = 200 μm (A) and 50 μm (C, D).

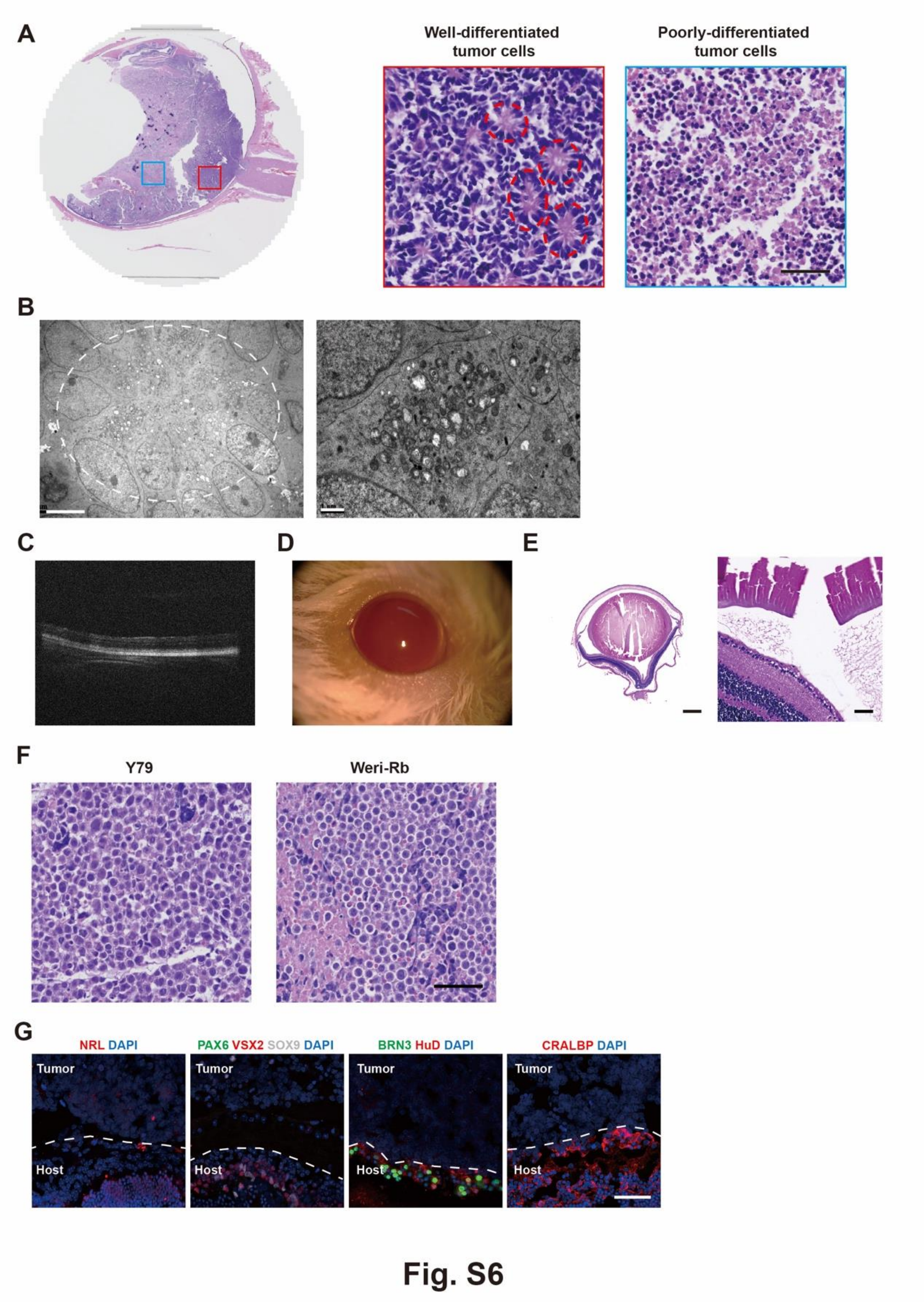


Fig. S6 Tumorigenesis of *RB1*$^{-/-}$ ROs *in vivo*

(A) Representative H&E staining images of Rb patient sample to represent the well-differentiated and poorly-differentiated tumor cells.

(B) Representative TEM image of tumor cells showing the formation of tumor after injection of cells from *RB1*$^{-/-}$ ROs with rosette structure the abundant mitochondria in 1st xenograft. Dash line represents the Flexner-Wintersteiner rosette structure.

(C) Representative OCT images of engrafted eye at 10 weeks after injection of WT.

(D) Representative slit-lamp images of engrafted eyes at 13 weeks after injection of WT. No tumor cells were observed in anterior chambers.

(E) Representative H&E staining images of retina after injection of cells from WT.

(F) Representative H&E staining images of eyes with injection of cells from Y79 or Weri-Rb-1 for 4 weeks.

(G) Representative immunostaining for NRL, PAX6, VSX2, BRN3, HuD and CRALBP in xenograft.

Scale bars = 50 μm (A, F, G), 5 (left) and 1 (right) μm (B), 500 (left) and 50 (right) μm (E).

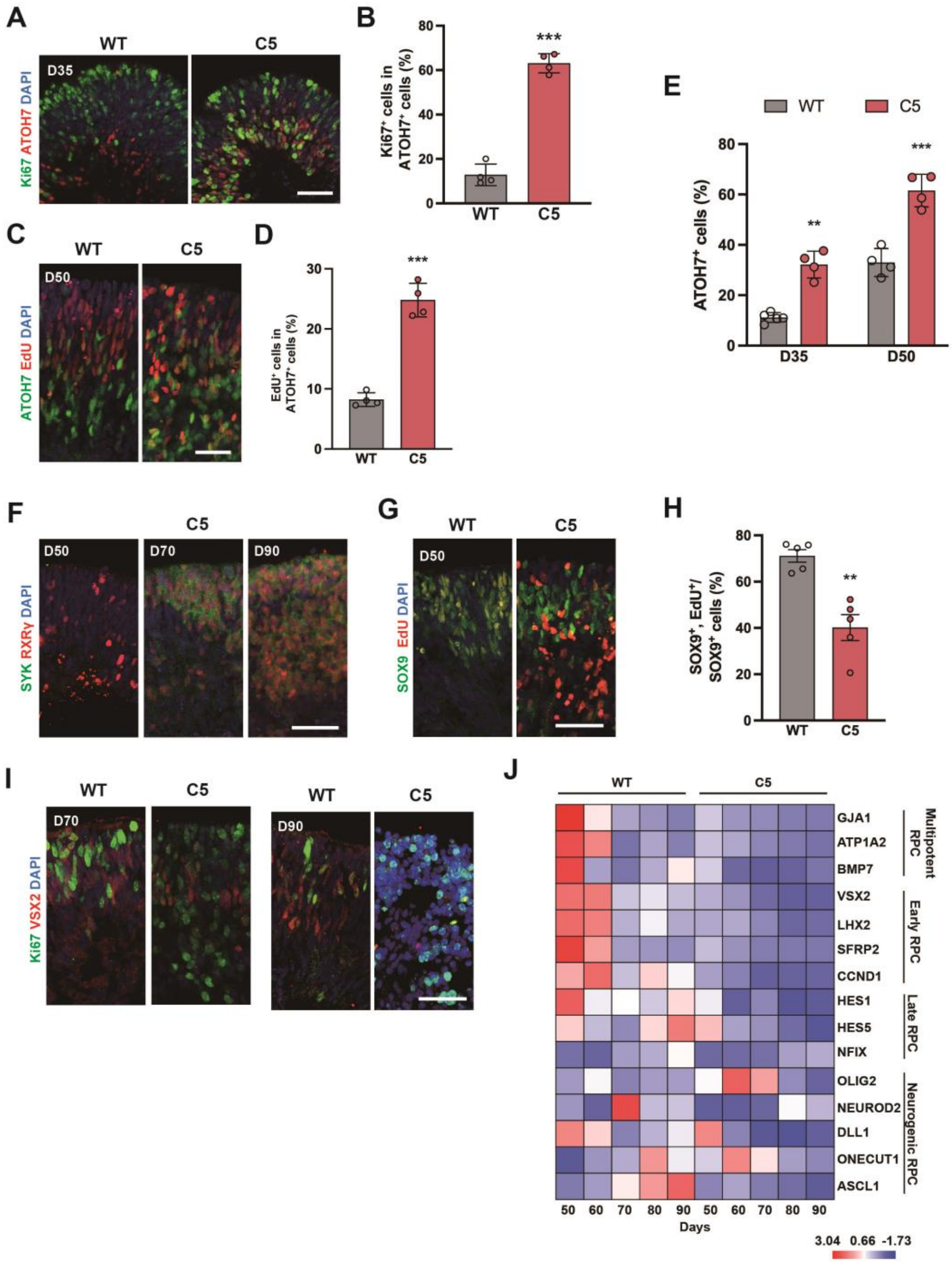


### Fig. S7

Fig. S7 Induction of cell cycle re-entry after *RB1* loss

(A) Representative immunostaining for Ki67 and ATOH7 in WT and *RB1*$^{-/-}$ ROs at day 35.

(B) Quantification of Ki67$^{+}$ cells in ATOH7$^{+}$ cells in WT and *RB1*$^{-/-}$ ROs at day 35. Data represent mean ± SD. *** P < 0.001 vs. WT, n = 4.

(C) Representative immunostaining for EdU and ATOH7 in WT and *RB1*$^{-/-}$ ROs after

EdU labeling.

(D) Quantification of EdU-labeled cells among ATOH7$^+$ in WT and *RB1*$^{-/-}$ ROs. Data represent mean ± SD. *** $P < 0.001$ vs. WT, n = 4.

(E) Quantification of the ratio of ATOH7$^+$ cells in WT and *RB1*$^{-/-}$ ROs at day 35 and 50. ** $P < 0.01$, *** $P < 0.001$ vs. WT, n = 4-5.

(F) Representative immunostaining for SYK and RXRγ in *RB1*$^{-/-}$ ROs from day 50 to 90.

(G) Representative immunostaining for EdU and SOX9 in WT and *RB1*$^{-/-}$ ROs after EdU labeling.

(H) Quantification of the ratio of EdU/SOX9$^+$ cells among SOX9$^+$ cells in WT and C5 *RB1*$^{-/-}$ ROs. Data represent mean ± SD. ** $P < 0.01$ vs. WT, n = 5.

(I) Representative immunostaining for Ki67 and VSX2 in WT and *RB1*$^{-/-}$ ROs from day 70 to 90.

(J) Heatmap of differential expression of RPC marker genes in WT and *RB1*$^{-/-}$ ROs from day 50 to 90.

Scale bars = 50 μm (A, C, F, G, I).

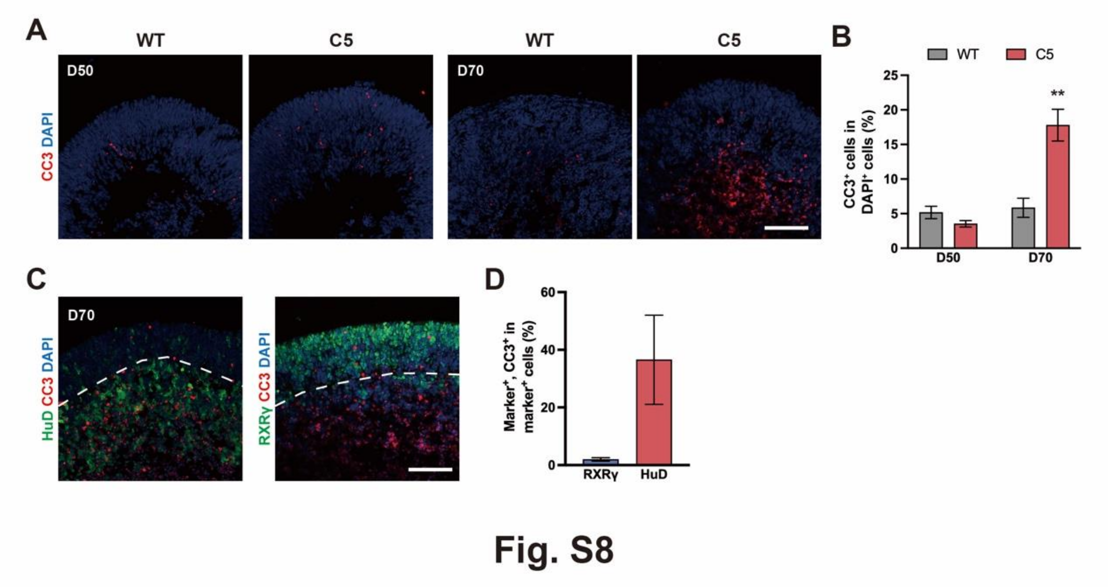


Fig. S8 Cell apoptosis in *RB1*$^{-/-}$ ROs-derived RGCs

(A) Representative immunostaining for cleaved caspase 3 (CC3) and HuD in WT and *RB1*$^{-/-}$ ROs at day 50 and day 70

(B) Quantification of $CC3^+$ cells in WT and *RB1*$^{-/-}$ ROs at day 50 and day 70. Data represent mean ± SD. ** $P < 0.01$ vs. WT, $n = 4$.

(C) Representative immunostaining for cleaved caspase 3 (CC3), HuD and RXRγ in *RB1*$^{-/-}$ ROs at day 70

(D) Quantification of the ratio of $CC3^+$ cells in $RXR\gamma^+$ cells or $HuD^+$ cells at day 70. Data represent mean ± SD, $n = 4$.

Scale bars = 100 µm (A, C).

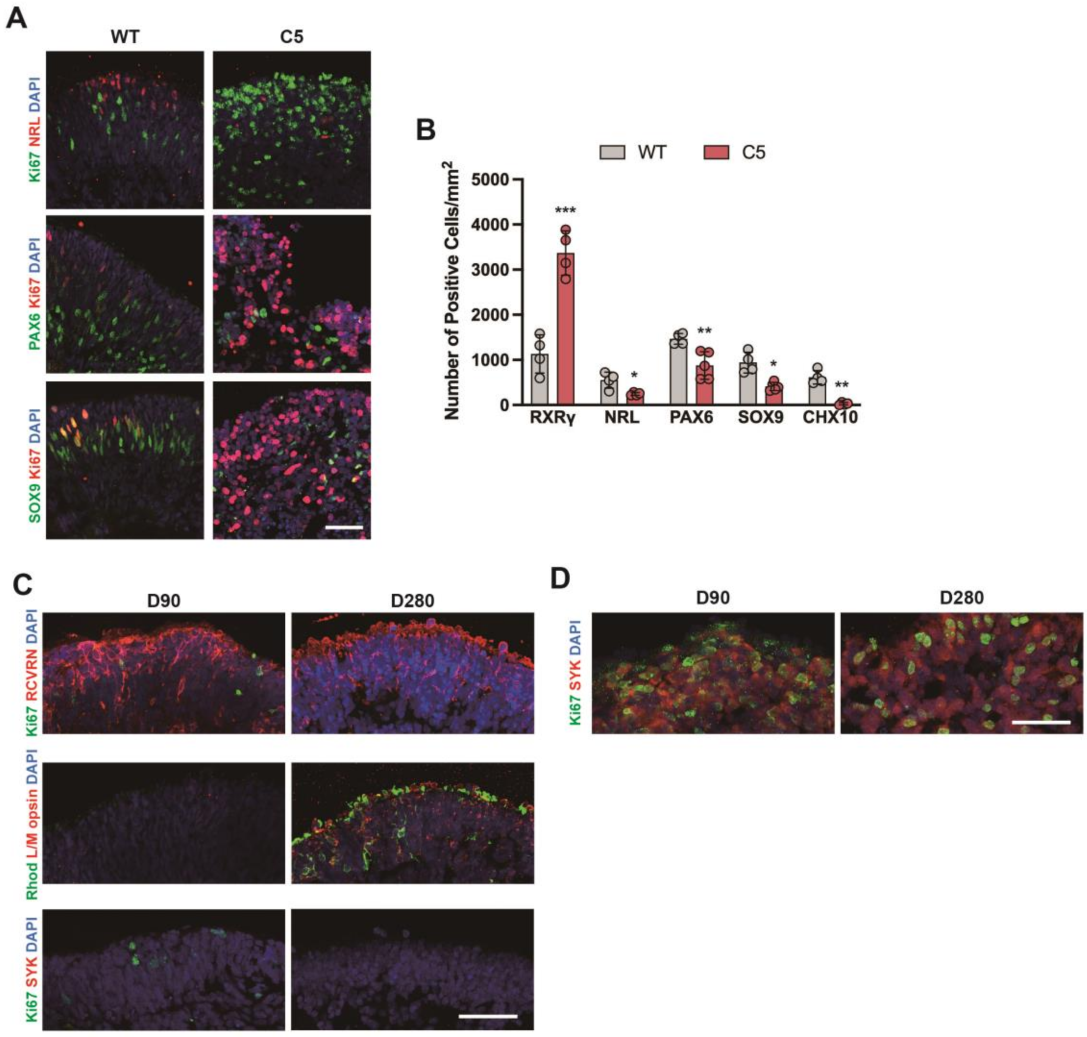


**Fig. S9**

Fig. S9 Maturation of *RB1*$^{-/-}$ ROs

(A) Representative immunostaining for NRL, PAX6, SOX9 and Ki67 in WT and C5 *RB1*$^{-/-}$ ROs at day 90.

(B) Quantification of cell number of RXRγ$^{+}$, NRL$^{+}$, SOX9$^{+}$, PAX6$^{+}$ and VSX2$^{+}$ cells in WT and *RB1*$^{-/-}$ ROs ROs at day 90. Data represent mean ± SD. * $P < 0.05$, ** $P < 0.01$, *** $P < 0.001$ vs. WT, n = 4-5.

(C) Representative immunostaining for RCVRN, Ki67, Rhodopsin, L/M opsin and SYK in WT ROs at day 90 and day 280.

(D) Representative immunostaining for Ki67 and SYK in *RB1*$^{-/-}$ ROs at day 160 and day 280.

Scale bars = 50 μm (A, C, D).

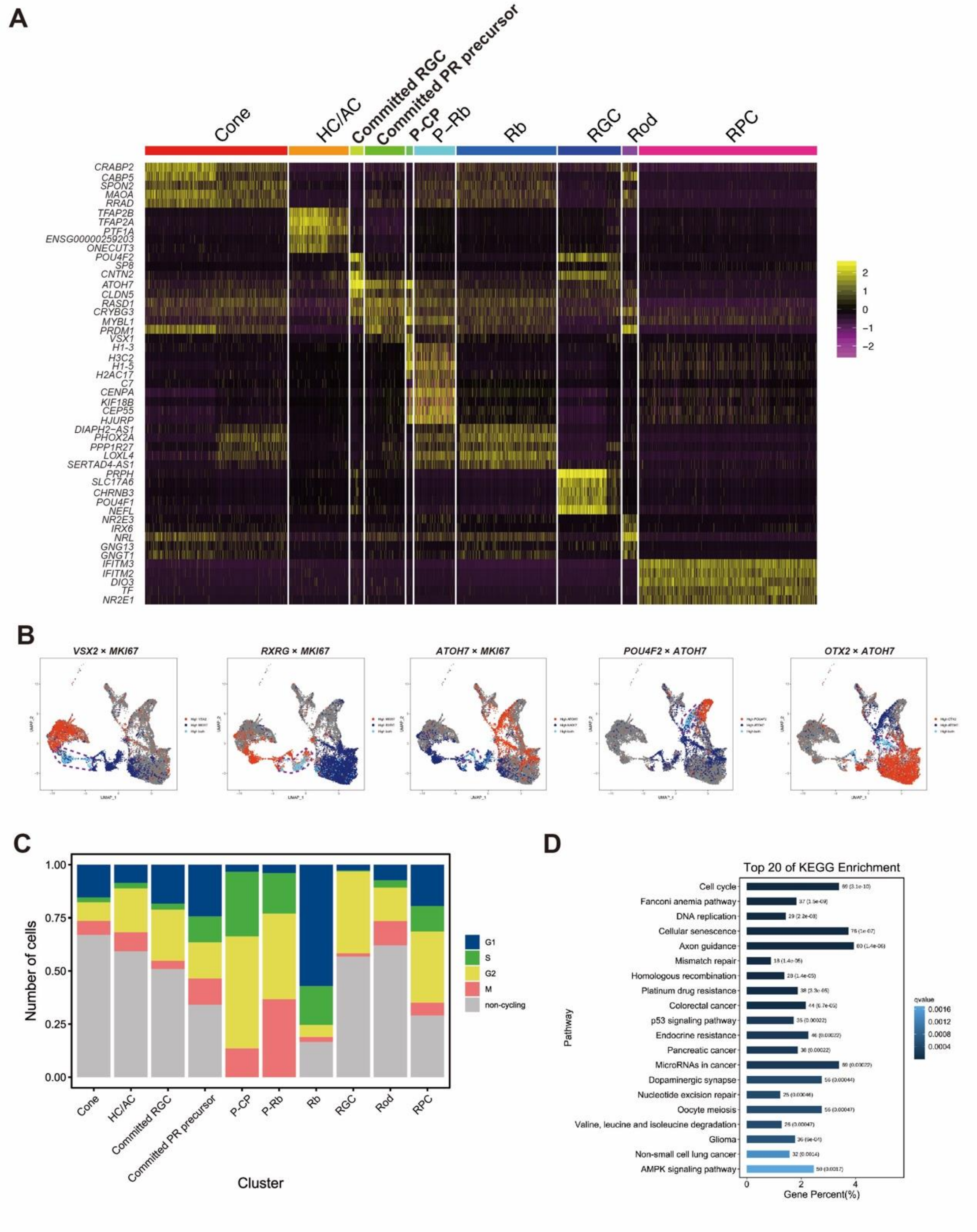


**Fig. S10**

Fig. S10 scRNA-seq of WT and *RB1*$^{-/-}$ ROs at day 80

(A) Heatmap visualization of enriched genes fir each cell types.

(B) Feature plots of double positive cells in WT and *RB1*$^{-/-}$ ROs.

(C) Quantification of ratio of cells in each cell cycle phase.

(D) KO enrichment for DEGs between Rb cells (Rb, P-Rb) and other retinal cell types.

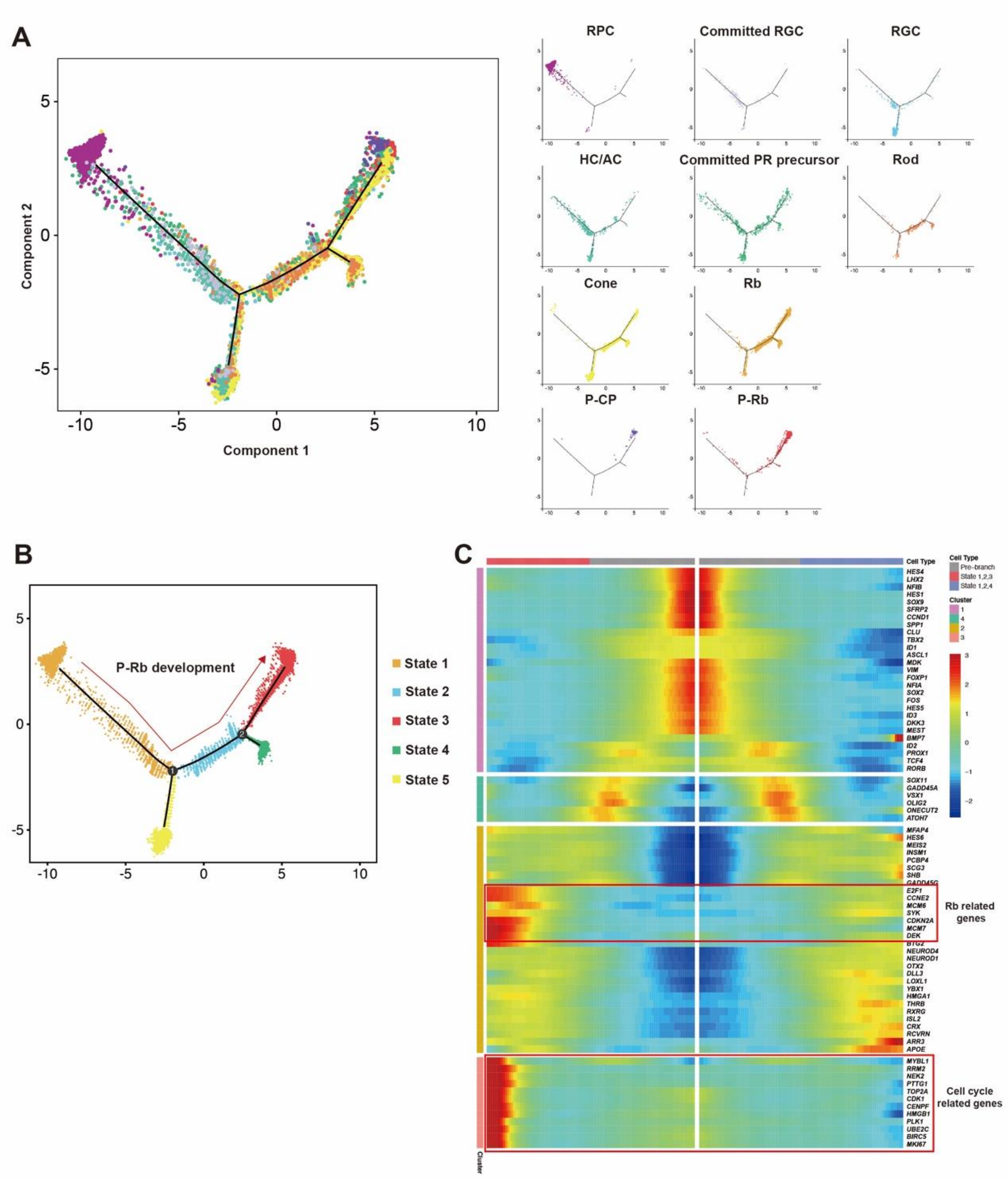


**Fig. S11**

<u>Fig. S11 Construction of Rb cells differentiation trajectory by *RB1*$^{-/-}$ ROs</u>

(A) Trajectory analysis of *RB1*$^{-/-}$ ROs by monocle 2 in each cell clusters.

(B) Pseudo-time trajectory analysis of *RB1*$^{-/-}$ ROs revealing the progress of P-CP into P-Rb cells.

(C) Heatmap showing the dynamic of selected Rb and cell cycle-related genes expression in P-Rb development.

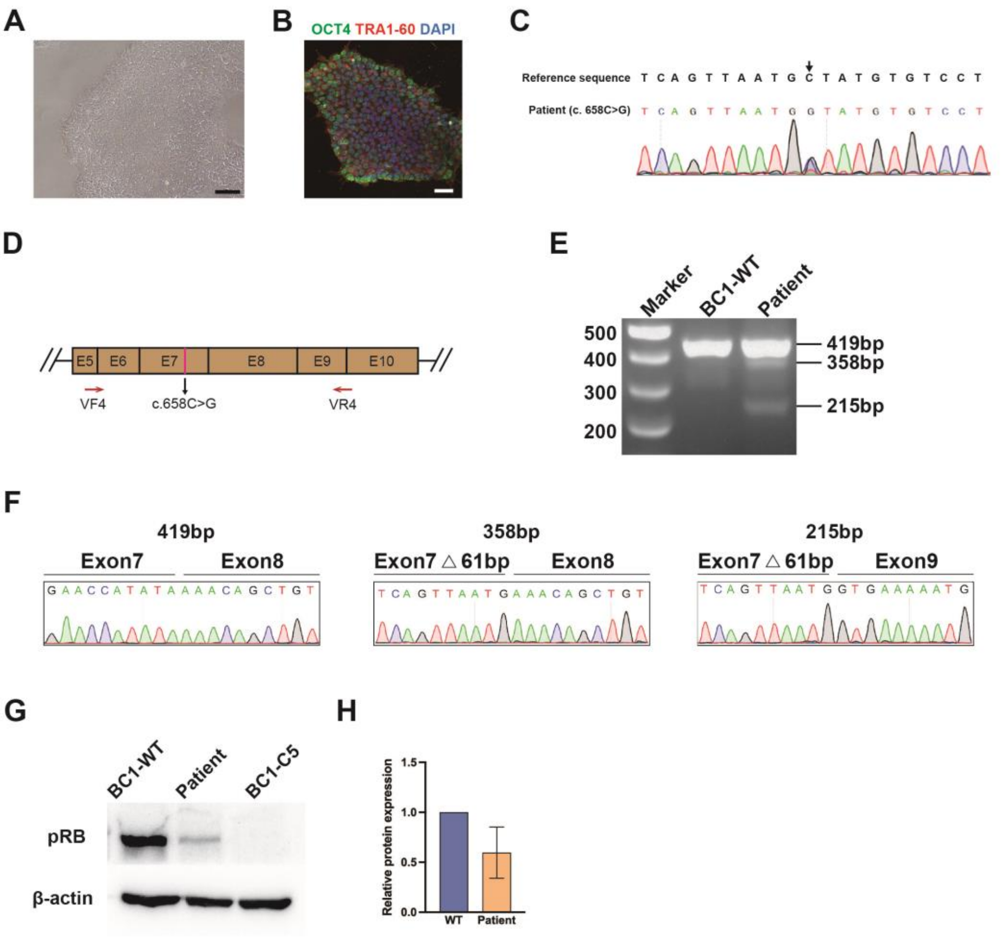


**Fig. S12**

<u>Fig. S12 Characterization of patient-specific hiPSC and ROs with *RB1* heterozygous mutation</u>

(A) Representative bright field images of patient-specific hiPSC line.

(B) Representative immunostaining images for OCT4 and TRA1-60 in patient-specific hiPSC line.

(C) Sequences of targeted loci in patient-specific hiPSC lines.

(D) Schematic diagram of the PCR primers used to analyze the effect of the c.658C>G mutation on mRNA splicing.

(E) RT-PCR amplification of the *RB1* transcripts obtained from patient-specific hiPSCs. Two types of additional bands were identified.

(F) Sequences corresponded to WT mRNA and Two types of alternatively spliced mRNA from patient-specific hiPSCs.

(G) Western blot analysis of pRB in WT, *RB1*$^{-/-}$ and patient-specific hiPSC line.

(H) Relative protein expression of pRB in WT hiPSC line and patient-specific hiPSC line. Data represent mean ± SD, n = 3.

Scale bars = 100 μm (B, C).

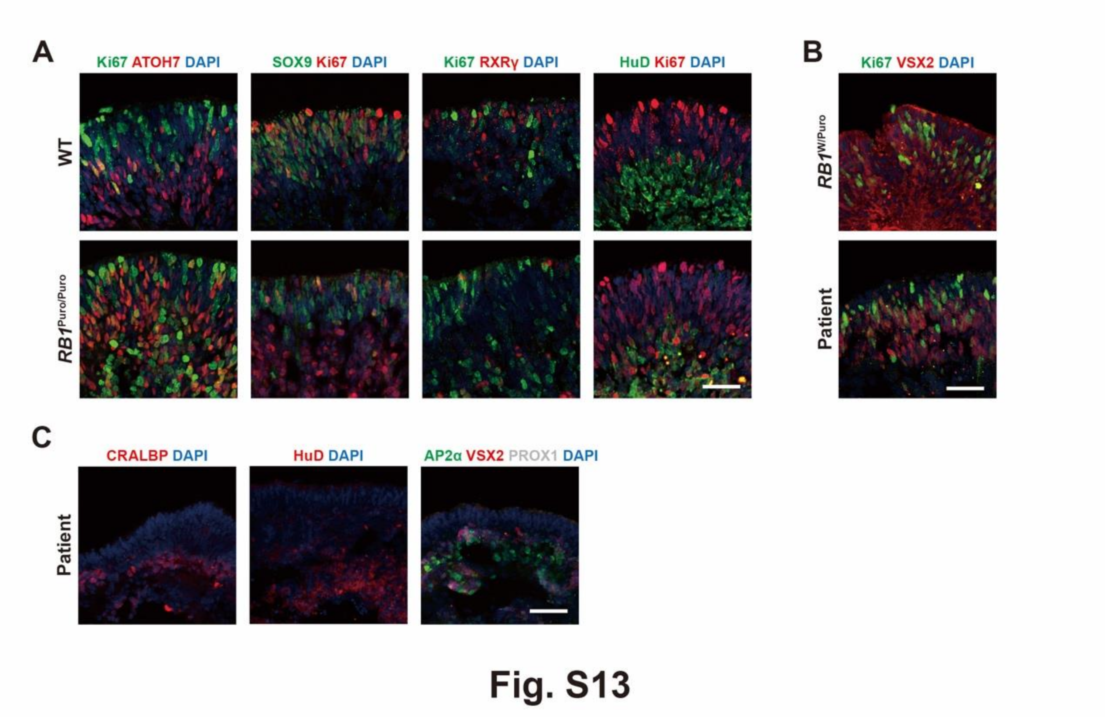


Fig. S13 Characterization of ROs with *RB1* heterozygous mutation

(A) Representative immunostaining for ATOH7, SOX9, RXRγ, HuD and Ki67 in WT and *RB1*$^{\text{Puro/Puro}}$ ROs (Gibco) at day 50.

(B) Representative immunostaining for RXRγ and Ki67 in patient-specific ROs (Gibco) at day 70 and 90.

(C) Representative immunostaining for CRALBP, HuD, AP2a, VSX2 and PROX1 in patient-specific ROs at day 160.

Scale bars = 50 μm (A, B, C).

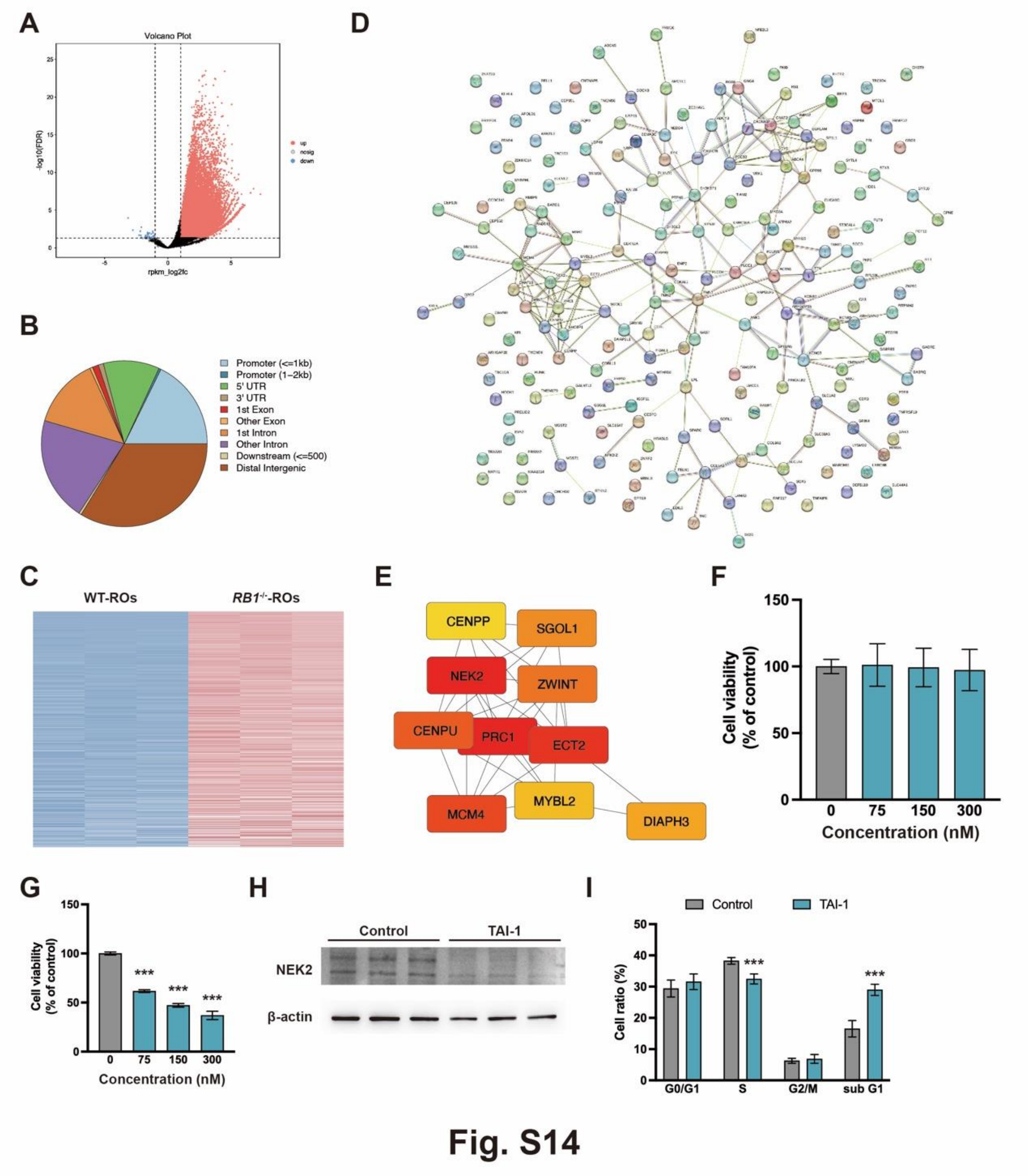


Fig. S14 Identification of NEK2 as a potential therapeutic target for Rb

(A) Volcano plot visualizations of differential peaks from ATAC-seq between WT and *RB1*$^{-/-}$ROs at day 80.

(B) Distribution of open chromatin across the whole genome in WT ROs

(C) Heatmap of the up-regulated genes between WT and *RB1*$^{-/-}$ ROs at day 80.

(D) Protein-protein interaction network of collective up-regulated genes from three data sets.

(E) Top ten hub genes from analysis of protein-protein interaction network by the cytoHubba plugin

(F) Cell viability of ARPE-19 after 72 hrs treatment with TAI-1 in different concentration by CCK8. Data represent mean ± SD, n = 6.

(G) Cell viability of Weri-Rb-1 cells after 72 hrs treatment with TAI-1 in different concentration by CCK8. Data represent mean ± SD. *** $P < 0.001$ vs. control, n = 6.

(H) Western blot analysis of NEK2 after TAI-1 treatment in Weri-Rb-1 cells.

(I) Cell cycle distribution of Weri-Rb-1 cells with TAI-1 treatment. Data represent mean ± SD. *** $P < 0.001$ vs. control, n = 6.

Supplementary Table 1 Summary of 1st xenograft using WT and *RB1*$^{-/-}$ ROs

| Cell types | Age of ROs | Number of transplanted cells | Total number of mice | Number of mice with tumor | Frequency |
|---|---|---|---|---|---|
| WT ROs | Day 61-85 | $1\times10^5$ | 5 | 0 | 0% |
| *RB1*$^{-/-}$ ROs | Day 67 | $1\times10^5$ | 10 | 7 | 70% |
| Y79 | N/A | $1\times10^5$ | 2 | 2 | 100% |
| Weri-Rb-1 | N/A | $1\times10^5$ | 2 | 2 | 100% |

Supplementary Table 2 Summary of 2nd xenograft using cells from 1st xenograft

| Cell types | Number of transplanted cells | Total number of mice | Number of mice with tumor | Frequency |
|---|---|---|---|---|
| *RB1*$^{-/-}$ ROs-derived 1st tumor | $1.2\times10^5$ – $1.5\times10^5$ | 5 | 5 | 100% |

Supplementary Table 3

| Oligonucleotides | |
|---|---|
| *RB1* sgRNA Targeting Sequence | GGTGGCGGCCGTTTTTCGGG |
| Primer VF1 | TTTGTAACGGGAGTCGGGAG |
| Primer VR1 | ACCTGTCAAGTTGAAGCCGA |
| Primer VF2 | CGGCCCTGGTATTGGACAAA |
| Primer VR2 | CAAGGGTAGCGGCGAAGAT |
| Primer VF3 | ACCATGCTGATAGTGATTGTTGAA |
| Primer VR3 | CCTGTCAGCCTTAGAACCATGT |
| Primer VF4 | ACACAACCCAGCAGTTCGAT |
| Primer VR4 | ACACAACCCAGCAGTTCGAT |

Supplementary Table 4 Information of antibody used in this research

| Antibodies | | |
|---|---|---|
| Goat polyclonal anti-ARR3 | Novus Biologicals | NBP1-37003 |
| Mouse monoclonal anti-AP2α (3B5) | Developmental Studies Hybridoma Bank | AB_528084 |
| Rabbit polyclonal anti-ATOH7 | Novus Biologicals | NBP1-88639 |
| Goat polyclonal anti-BRN3 (C13) | Santa Cruz | sc-6026 |
| Rabbit polyclonal anti-Cleaved Caspase 3 | Cell Signaling Technology | #9661 |
| Rabbit polyclonal anti-CDKN2A/p16$^{INK4a}$ | Bioss | bs-20656R |
| Mouse monoclonal anti-CRX(M02) Clone 4G11 | Abnova | H00001406-M02 |
| Mouse monoclonal anti-HuD(H-9) | Santa Cruz | sc-48421 |
| Mouse monoclonal anti-Ki67(B56) | BD Biosciences | 550609 |
| Rabbit polyclonal anti-Ki67 | Abclonal | A11390 |
| Rat monoclonal anti-Ki67 | Origene | TA801577S |
| Rabbit anti-L/M opsin | Gift from Dr Jeremy Nathans | N/A |
| Rabbit polyclonal anti-NANOG | Abcam | ab21624 |
| Mouse monoclonal anti-TRA-1-60 | Abcam | ab16288 |
| Mouse monoclonal anti-SSEA4 (MC813-70) | Abcam | ab16287 |
| Rabbit polyclonal anti-OCT4 | Abcam | ab19857 |
| Rabbit polyclonal anti-NEK2 | Proteintech | 14233-1-AP |
| Mouse monoclonal anti-NR2E3 | R&D Systems | PP-H7223-00 |
| Mouse monoclonal anti-NRL (F-2) | Santa Cruz | sc-374277 |
| Goat polyclonal anti-NRL | R&D Systems | AF2945 |
| Rabbit polyclonal anti-OTX1+OTX2 | Abcam | ab21990 |
| Mouse polyclonal anti-PAX6 | DSHB | AB_528427 |
| Rabbit polyclonal anti-PROX1 | Millipore | AB5475 |
| Rabbit monoclonal anti-Rb | Abcam | ab181616 |
| Mouse monoclonal anti-Rb (G3-245) | BD Biosciences | 554136 |
| Mouse monoclonal anti-Rb (4H1) | Cell Signaling Technology | #0309 |

| | | |
|---|---|---|
| Rabbit polyclonal anti-RCVRN | Millipore | AB5585 |
| Mouse monoclonal anti-Rhodopsin (1D4) | Abcam | ab5417 |
| Rabbit polyclonal anti-RXRγ | Abcam | ab15518 |
| Mouse monoclonal anti-RXRγ (A-2) | Santa Cruz | sc-365252 |
| Rabbit monoclonal anti-SOX9 (ARC0190) | Abclonal | A19710 |
| Mouse monoclonal anti-STEM121 | Cellartis(Takara) | Y40410 |
| Rabbit monoclonal anti-SYK (D3Z1E) | Cell Signaling Technology | #13198 |
| Mouse monoclonal anti-SYK (4D10) | Santa Cruz | sc-1240 |
| Rabbit polyclonal anti-α-Tubulin | Beyotime | AF0001 |
| Sheep polyclonal anti-VSX2 | Millipore | AB9016 |
| Donkey Anti-Mouse IgG-488 | Invitrogen | A21202 |
| Donkey Anti-Rabbit IgG-488 | Invitrogen | A21206 |
| Donkey Anti-Rat IgG-488 | Invitrogen | A21208 |
| Donkey Anti-Mouse IgG-555 | Invitrogen | A31570 |
| Donkey Anti-Rabbit IgG-555 | Invitrogen | A31572 |
| Donkey Anti-Goat IgG-555 | Invitrogen | A21432 |
| Donkey Anti-Sheep IgG-555 | Invitrogen | A21436 |
| Donkey Anti-Rabbit IgG-647 | Invitrogen | A31573 |
| Donkey Anti-Mouse IgG-647 | Invitrogen | A31571 |
